# Chromosome-scale genome assembly of the brown anole (*Anolis sagrei*), a model species for evolution and ecology

**DOI:** 10.1101/2021.09.28.462146

**Authors:** Anthony J. Geneva, Sungdae Park, Dan Bock, Pietro de Mello, Fatih Sarigol, Marc Tollis, Colin Donihue, R. Graham Reynolds, Nathalie Feiner, Ashley M. Rasys, James D. Lauderdale, Sergio G. Minchey, Aaron J. Alcala, Carlos R. Infante, Jason J. Kolbe, Dolph Schluter, Douglas B. Menke, Jonathan B. Losos

## Abstract

Rapid technological improvements are democratizing access to high quality, chromosome-scale genome assemblies. No longer the domain of only the most highly studied model organisms, now non-traditional and emerging model species can be genome-enabled using a combination of sequencing technologies and assembly software. Consequently, old ideas built on sparse sampling across the tree of life have recently been amended in the face of genomic data drawn from a growing number of high-quality reference genomes. Arguably the most valuable are those long-studied species for which much is already known about their biology; what many term emerging model species. Here, we report a new, highly complete chromosome-scale genome assembly for the brown anole, *Anolis sagrei* – a lizard species widely studied across a variety of disciplines and for which a high-quality reference genome was long overdue.

## Introduction

Recent breakthroughs in high-throughput sequencing, coupled with the creation of long-distance scaffolding libraries, have ushered in an era of ever-improving quality and quantity of genome assemblies. Genome assemblies now routinely span entire chromosomes and include data from formerly impenetrable genomic regions^1-3^. In turn, these assemblies have enabled increasingly sophisticated genomic analyses of organismal traits and behaviors, and the evolutionary and ecological implications of the interactions of genomes and the environment. Massive reductions in the cost of genome sequencing and assembly have allowed non-model and emerging model species to become genome-enabled, a neologism indicating that genomic information has become available for the species. Observations from these new assemblies have provided fresh insights into core biological processes. For example, our understanding of recombination^4^, repetitive genetic elements^5,6^, chromosome evolution^7-9^ and dosage compensation^4,10,11^ have all been fundamentally amended due to results made possible by recent genome assemblies of non-traditional model species.

Efforts are underway to generate thousands of new genome assemblies for species across the tree of life^12-14^. However, our understanding of the biology of most species on earth remains sorely lacking – limiting the inferential power gained by the addition of genomic data. In contrast, those species for which the existing organismal literature is vast are particularly primed for the generation of new, high-quality genome assemblies because new discoveries concerning the genetic basis of organismal traits await only the addition of a highly complete and contiguous reference genome.

While the production of highly contiguous genome assemblies is a technological achievement, the long-term value of these assemblies is that they serve as critical tools in the advancement of biological research. Evolutionary genomic techniques such as quantitative trait locus mapping or genome-wide association studies enable careful examination of the genetic basis of organismal traits, but these rely on linkage disequilibrium information to connect genotype to phenotype. Improved contiguity of genome assemblies therefore paved the way for a finer and more accurate understanding of the genomic basis of organismal traits.

Further, understanding the evolutionary history of a species’ chromosomes similarly requires highly complete genome assemblies since only with these data can chromosomal sequence homology be reliably inferred^15,16^. While cytogenetics opened the door to inferring evolutionary transitions in chromosome complement well over 100 years ago^17^, only recently through genome sequencing have the evolutionary drivers and consequences of these changes begun to be understood. While the first wave of genome assemblies lacked the contiguity and completeness to fully determine syntenic relationships between species, new chromosome-scale assemblies now enable rigorous study of chromosome evolution.

Finally, population genomic scans also benefit from improved contiguity. For example, recent selective sweeps leave patterns of reduced genetic diversity in the genomic regions surrounding the selected variant. Many methods to detect recent selection rely on these patterns but poorly constructed genome assemblies can separate that signal onto separate scaffolds and limit our ability to detect these patterns.

### The Brown Anole

*Anolis* lizards (anoles) comprise over 400 small-to medium-sized lizard species distributed throughout the continental neotropics of South, Central, and North America, and across islands in the West Indies and eastern Pacific Ocean^18^. The green anole (*Anolis carolinensis*) was the first reptile to have its full genome assembled^19^. While it was sequenced using first-generation genome sequencing technologies over 10 years ago, it remains one of the best assembled and annotated reptile genomes and by far the most complete and contiguous assembly within the genus *Anolis*. It was selected for genome sequencing due to many decades of biomedical research–especially epidemiology and neurobiology–using this species as a model. Recently, a second species, the brown anole (*Anolis sagrei*), has surpassed the green anole in the number of publications per year (Fig. 1) and is considered an emerging model species for numerous fields.

**Figure 1.**
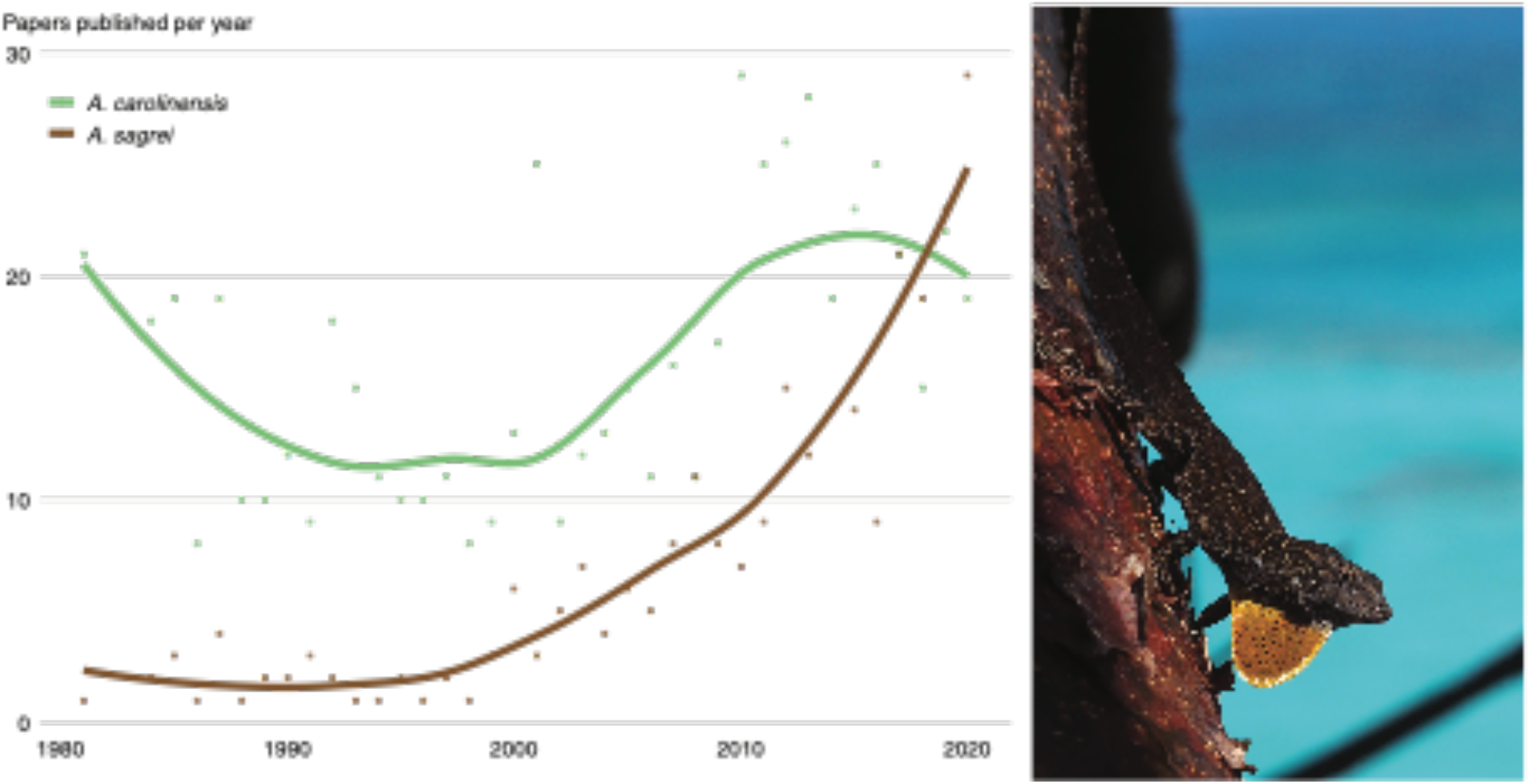
The rise of *Anolis sagrei*. Over the past 40 years research interest in *Anolis sagrei* (pictured at right) has grown substantially and recently surpassed that of *A. carolinensis*, which for many decades served as the model reptile species in biological research. Queries for each specific epithet were performed in the indexed Titles and Abstracts on https://www.dimensions.ai (accessed May 2021).

*Anolis sagrei* is a medium-sized insectivorous lizard most commonly found on the ground and perched low on the trunks of trees^20^. Although it first arose on Cuba, the species now has the largest native range of any anole with natural diaspora populations found across islands of the northern Caribbean as well as coastal areas of Mesoamerica^21,22^. It is also a prolific invader with non-native populations established on many additional islands in the West Indies^23,24^, Costa Rica, multiple locations in both North^25^ and South America^26,27^, as well as remote islands of the central Atlantic Ocean^28,29^, Hawaii^30^, Taiwan, and mainland Asia, Europe, and the Middle East.

A recent analysis of genome-scale sequence data revealed that *A. sagrei* evolved on Cuba toward the end of the Miocene^22^. Two major lineages are present on East and West Cuba, and although they are not geographically separate, they represent ancient evolutionary separation and probable recent secondary contact. Both lineages have given rise to diaspora populations that have colonized other island groups. The western Cuba lineage colonized the Bahamas Archipelago in both the Pliocene and Pleistocene, while the eastern lineage colonized the Cayman Archipelago, the Swan Islands, Mesoamerica, and Jamaica at different periods during the Pleistocene^22^. These diaspora lineages, despite different evolutionary backgrounds and divergence times, have evolved a similar suite of phenotypic traits such that Cuban *A. sagrei* can be distinguished from diaspora *A. sagrei* using both genetic and phenotypic characters^22^. This suggests that the species has responded to presumably similar evolutionary selective pressures when colonizing islands elsewhere in the Caribbean. Notably, both relatively larger body size and increased number of subdigital lamellar scales appear to be features of diasporic lineages, although it is currently unknown whether similar genomic changes are responsible for these outcomes.

Multiple factors have led to the rapidly increasing use of *A. sagrei* for research in evolution and ecology. These include its wide natural and invasive ranges, its high local abundance, and the fact that this species is amenable to captive treatments including breeding and rearing in a laboratory setting^31,32^. As a result, this species has been the focus of years of detailed evolutionary, developmental, ecological, behavioral, and physiological research conducted both in natural environments and in the lab^33^. Over the past three decades, the brown anole has become a broadly used system to study evolutionary ecology^34-36^, behavior^37,38^, development^39-43^, reproductive isolation^44^, sexual selection^45-47^, biological invasions^48-51^, and adaptation^52-54^. However, the lack of a reference genome has made it challenging to connect this depth of knowledge of brown anole phenotypes to their underlying genetic architecture. Despite this limitation, the brown anole has been at the forefront of new techniques including chromosome microdissection and sequencing^55,56^ and recently became the first reptile to successfully undergo CRISPR-Cas9 genome editing^57^. This last breakthrough begs for the production of a high-quality reference genome to establish the brown anole as a fully-fledged model organism.

Here, we report a highly complete and contiguous genome assembly of a single female brown anole (*Anolis sagrei ordinatus*) from the Central Bahamas. We supplement this assembly with evidence-based and *ab initio* gene model annotation, repetitive element identification and analysis, and a map of segregating genetic diversity. Finally, we build on existing research to confirm the identity of the *A. sagrei* X chromosome and identify patterns of the evolution of the *A. sagrei* X chromosome relative to its counterparts in the *A. carolinensis* genome.

## Results and Discussion

We created a highly complete and contiguous draft genome assembly of *A. sagrei* through multiple rounds of iterative improvement. Our initial assembly using only Illumina whole-genome shotgun sequences and assembled using meraculous^58^ produced a largely fragmented assembly, which was incomplete in terms of gene content and total size (Table S1). Subsequent scaffolding performed in HiRise^59^ using Chicago and HiC proximity ligation libraries substantially improved both contiguity and completeness, but the assembly remained substantially smaller (1.6Gb) than the 1.8Gb assembly of *A. carolinensis* and a genome size estimate of 1.89Gb for *A. sagrei* based on fluorescence cytophotometry^60^. We further refined the *A. sagrei* genome assembly by improving contig size with error-corrected PacBio long reads and re-scaffolding in HiRise. The addition of these data resulted in a far more contiguous and complete assembly, the size of which (1.93 Gb) very closely matches the expected genome size for this species (Table S1). Analysis of HiC mapped read link density using Juicer v1.6^61^ revealed that two chromosomes had been artificially joined during the assembly process. Using evidence from Illumina short-read, RNA-Seq, and PacBio data (see Methods) we corrected this misjoin resulting in the current *A. sagrei* assembly version (hereafter, AnoSag2.1). A link density histogram of HiC read pairs mapped to the AnoSag2.1 assembly does not show evidence of remaining misjoins (Fig. 2c). The mitochondrial genome was not captured in this assembly but was recovered through a combination of circularized *de novo* assembly and identification of mitochondrial sequence in an error-corrected PacBio read. The consensus of these two approaches yields a 17,535bp assembly with the 13 genes, 22 tRNAs, and two ribosomal RNAs expected for vertebrates and with identical gene ordering to the *A. carolinensis* mitochondrial genome.

**Figure 2.**
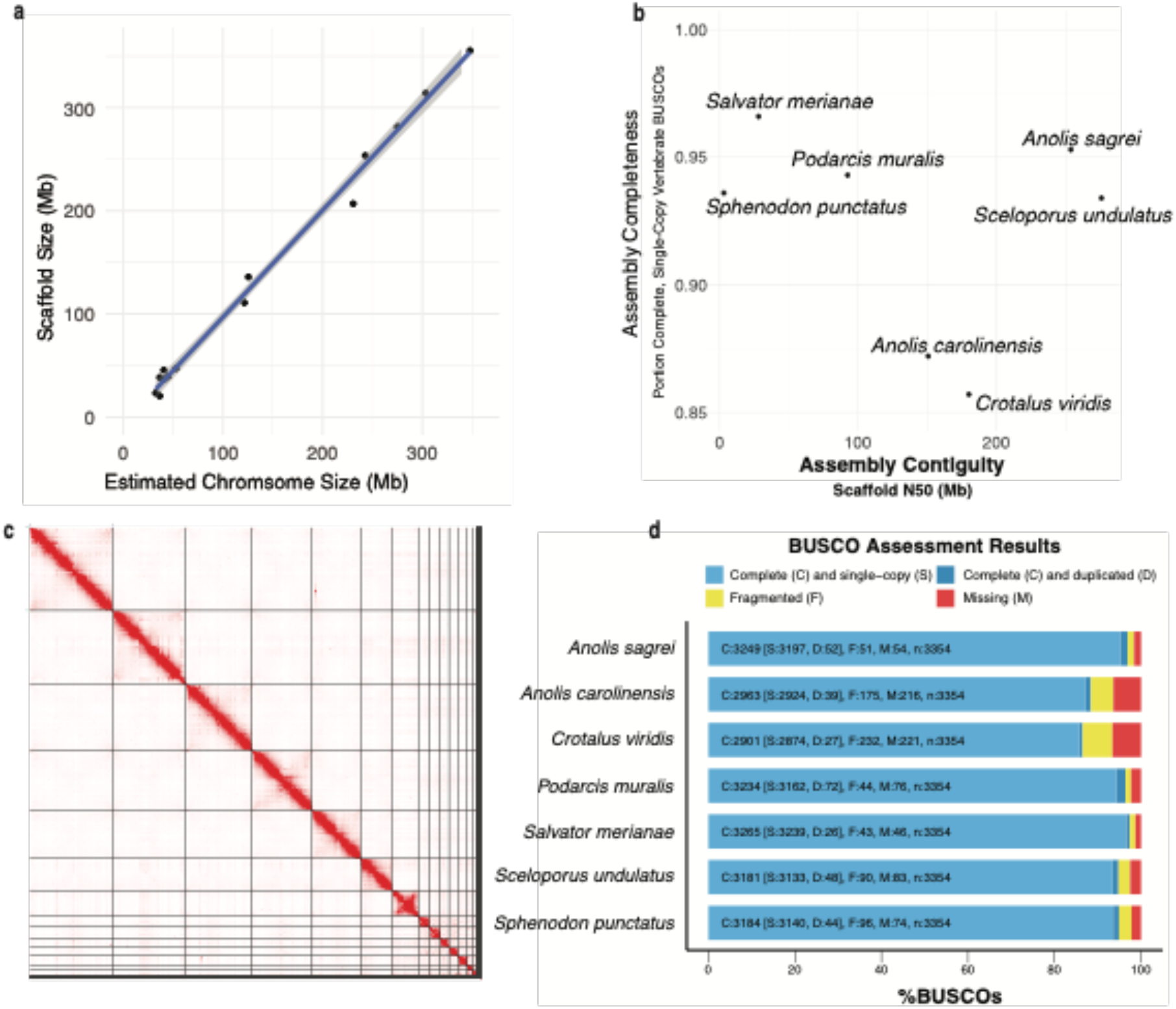
Contiguity and completeness of *Anolis sagrei* and other lepidosaur assemblies. **a**) The scaffold sizes of the AnoSag2.1 assembly are highly correlated with chromosome sizes estimated from karyotype imaging. **b**) Scatterplot of recent lepidosaur gene assemblies **c**) Link density histogram of the AnoSag2.1 assembly **d**) BUSCO assessment of assembly completeness for AnoSag2.1 and other lepidosaur assemblies.

### Contiguity and Completeness

Our AnoSag2.1 assembly has a scaffold N50 of 253.6Mb, which is 1.6 times as contiguous as *Anolis carolinensis*, the longtime standard bearer for reptile genome assemblies^19^. The four largest AnoSag2.1 scaffolds comprise more than 50% of the genome assembly. The *A. sagrei* karyotype contains 14 chromosomes: six macrochromosomes, seven microchromosomes and the intermediately sized X chromosome. Multiple lines of evidence suggest that our assembly recovers each of these chromosomes as the 14 largest scaffolds. First, the 14 largest scaffolds in AnoSag2.1 comprise 99.1% of the assembled genome sequence. Furthermore, a large drop-off in scaffold size occurs after the last putative chromosome – scaffold 14 is over 20Mb in size where the next largest scaffold is two orders of magnitude smaller (scaffold 15; 131kb). Finally, the AnoSag2.1 scaffold sizes are highly correlated (r2=0.996, p< 2.2x e-16) with chromosome sizes estimated using a published karyotype^62^ of this species (Fig. 2a).

We assessed completeness of our assembly using BUSCO 5.0.0 which tests for the presence of a curated set of 3,354 protein-coding genes known to be present in single copy across vertebrate genomes (vertebrata_odb10). Of these genes, 3,197 (95.3%) are present in full length and found to be single-copy in our assembly. The AnoSag2.1 assembly is missing only 1.6% of the genes from this set. Our assembly exceeds most other Lepidosauria (lizards, snakes, and tuatara) genome assemblies in contiguity and completeness^4,19,63-66^ (Table S2). Only the Argentine black and white tegu^66^ (*Salvator merianae*) exceeds our assembly in BUSCO completeness but is substantially less contiguous (Fig. 2b). The eastern fence lizard^64^ (*Sceloporus undulatus*) is slightly more contiguous than our assembly but less complete. These two genomes stand apart from other recent lepidosaur genome assemblies in being both highly complete and contiguous (Fig. 2b,d).

### Annotation Statistics

We performed an automated annotation of our assembly using Braker v2.0.5^67^ followed by manual curation for roughly 15% of all gene models. This effort resulted in a final set of 21,853 genes comprising 849 Mb (44.1% of the final assembly length). Most gene models (92%) contain more than one exon and all exons summed account for a total length of 55 Mb, or about 3% of the assembly. Start codons are annotated for 99.7% of all gene model and the same percentage have stop codons annotated (although not all within the same genes). BUSCO analysis of the annotated exome suggests our annotation captures most of the genes found in the homology-based BUSCO search – 95% of vertebrate universal single-copy orthologs were found to complete and single copy via homology search of the entire genome sequence versus 90% found within the gene models in our annotation (Table S3).

### Repetitive Element Landscape

We estimated 46.3% of the *Anolis sagrei* genome to be repetitive, compared with 32.9% for *A. carolinensis*. Both genomes contained a diversity of transposable elements, including short interspersed elements (SINEs), long interspersed nuclear elements (LINEs), long terminal repeat retrotransposons (LTRs), and DNA transposons (Table 1). *Anolis sagrei* contained a higher proportion of LINEs and DNA transposons, whereas *A. carolinensis* contained relatively more LTR retrotransposons.

**Table 1.** Repetitive Elements. Comparison of the interspersed repeat contents of *Anolis sagrei* and *Anolis carolinensis*.

| Repeat Class | *Anolis sagrei* |  | *Anolis carolinensis* |  |
| --- | --- | --- | --- | --- |
|  | Length occupied (bp) | Percent of Genome | Length occupied (bp) | Percent of Genome |
| SINEs | 71,051,564 | 4.48 | 75,887,012 | 4.22 |
| LINEs | 308,353,200 | 19.44 | 234,058,101 | 13.01 |
| LTR elements | 28,262,816 | 1.78 | 84,049,288 | 4.67 |
| DNA transposons | 175,148,224 | 11.04 | 157,677,814 | 8.76 |
| Unclassified | 137,820,970 | 8.69 | 34,170,372 | 1.90 |
| **Total** | **720,636,774** | **45.43** | **585,842,587** | **32.56** |

We examined the age distribution of repeats in each genome, or its repeat landscape, by comparing the proportion of the assembly comprised of insertions according to their divergence from family consensus. When comparing the repeat landscapes of the anole genomes, we found that *A. carolinensis* contained a much higher proportion of transposable element insertions with ≤10% divergence from their family consensus (Fig. 3). This was for DNA transposons (Kruskal Wallis test; P=0.0005), LTR retrotransposons (P=8.09e-07), and LINEs (P=0.0073), but not SINEs. This suggests that while the transposable element landscape of the *A. sagrei* genome includes more DNA transposons and LINEs than *A. carolinensis*, this discrepancy is driven by a much larger proportion of the genome comprised of ancient insertions beyond 10% Kimura 2-parameter divergence in *A. sagrei*. In contrast, the transposable element landscape of the *A. carolinensis* genome is dominated by recent inserts which is indicative of recent activity.

**Figure 3.**
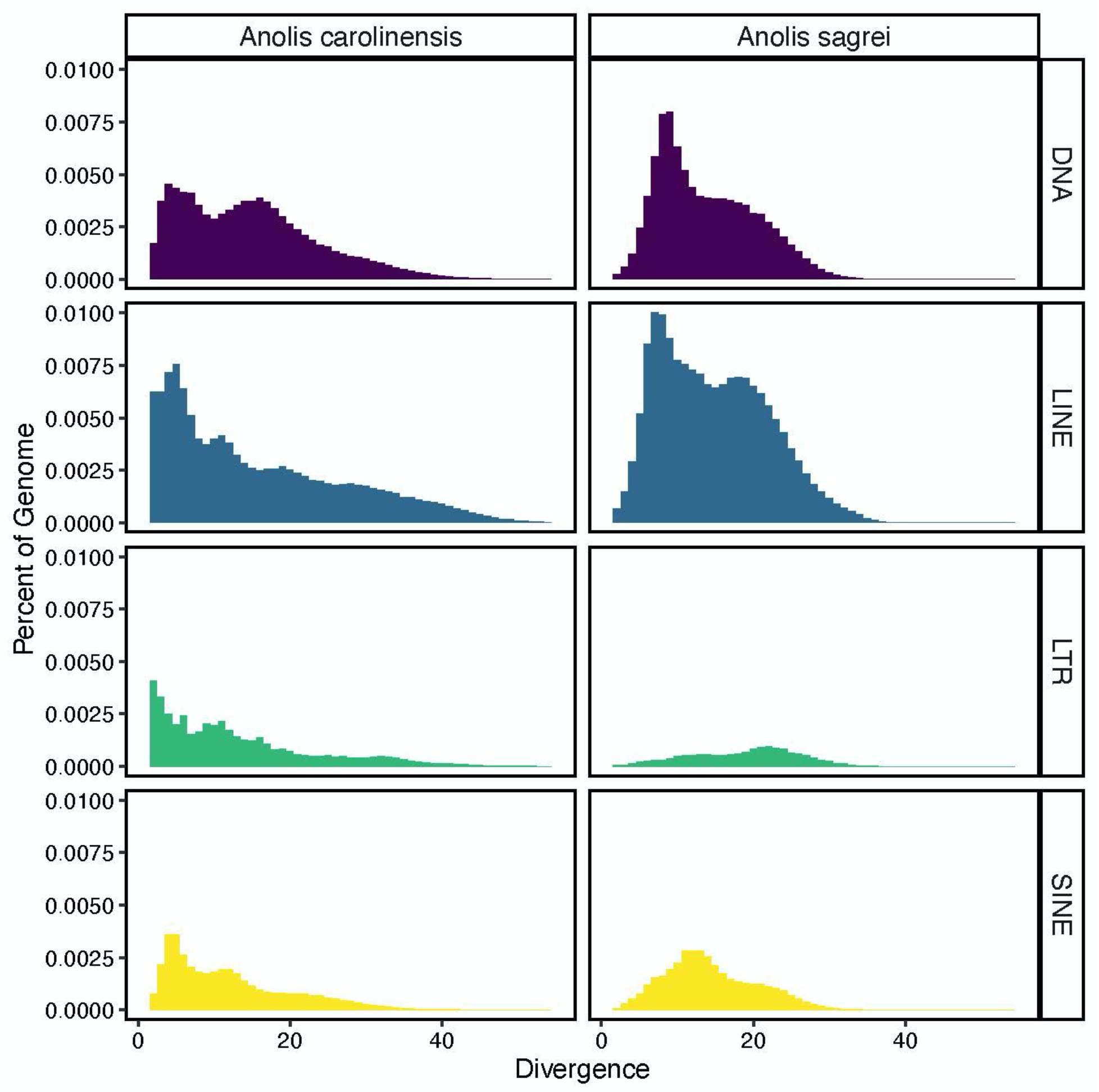
Comparison of repeat landscapes for the classes of transposable elements in *Anolis carolinensis* and *Anolis sagrei*. The proportion of the genome consisting of transposable element insertions (short interspersed elements=SINE, long interspersed elements=LINE, long terminal repeat retrotransposons=LTR, and DNA transposons=DNA) of different ages according to their Kimura 2-parameter divergence from consensus. Older insertions are more divergent.

### Synteny and Sex Chromosome analyses

A growing body of evidence suggests that fusions between autosomes and sex chromosomes are common across anoles^68,69^. The chromosomes that result from such fusions, called neo-sex chromosomes, have been used to study multiple evolutionary processes, such as chromosomal degeneration^70^ (or lack thereof) and dosage compensation^71^. *Anolis sagrei* has an XY sex determination system with sex chromosomes that are significantly larger than those from *A. carolinensis*^*62,68*^. As Iguanian lizards (with the exception of Basilisks) share a highly conserved core of X-linked genes^72^, the enlarged sex chromosomes in *A. sagrei* have been hypothesized to be the product of three independent fusions of autosomes to conserved iguanian X and Y sex chromosomes^55,73^. By aligning *A. sagrei* short reads from chromosomal flow sorting to the reference *A. carolinensis* genome, Giovannotti and colleagues^62^ showed that the 7th largest chromosome in the *A. sagrei* karyotype was the product of a series of chromosomal fusion events that occurred in the *A. sagrei* lineage. Ancestral chromosomes homologous to *A. carolinensis*’ chromosomes 9 and 12 fused to chromosome 13, the X chromosome of *A. carolinensis* (henceforth ‘ancient X’). Soon after, Kichigin and colleagues found that the chromosome corresponding to *A. carolinensis* chromosome 18 had also fused to the ancient X in the *A. sagrei* lineage^56^. These authors hypothesized that the neo-sex chromosome in *A. sagrei* resulted from three fusion events: chromosomes 12 and ancient X would have fused independently from chromosomes 9 and 18, and these two pairs of fused chromosomes then fused together to create the current *A. sagrei* XY system. Kichigan further proposed a synteny hypothesis for the *A. sagrei* neo-sex chromosomes in which the ancient X and chromosome 18 would be at the extremes of the neo-X chromosome, while chromosomes 12 and 9 would be in its center^56^.

Analyses of read depth, heterozygosity, and genome-wide association all indicate that AnoSag2.1 scaffold 7 is the X chromosome in this species (see *Sex Chromosome Identification* below). Using SatsumaSynteny, we aligned the *A. carolinensis* and *A. sagrei* genomes and confirmed previously published predictions^56,73^ that the X chromosome in *A. sagrei* is the product of fusions between chromosomes homologous to 9, 12 and 18 from the *A. carolinensis* assembly and the ancient X. Given the level of contiguity of the AnoSag2.1 scaffold 7, our results present a clear synteny prediction for not only the order of the *A. carolinensis* chromosomes in scaffold 7, but also for the linkage groups that make up the ancient X in *A. carolinensis* (Fig. 4g). We found extensive overlap between the list of scaffolds identified in our synteny analyses and those obtained using short-read data from chromosomal flow sorting^73^. Furthermore, our data corroborated previous results based on dosage compensation, qPCR of X-linked genes and flow sorting in *A. carolinensis* that identified 8 additional scaffolds as X-linked in the original *A. carolinensis* assembly^11,56,74,75^.

**Figure 4.**
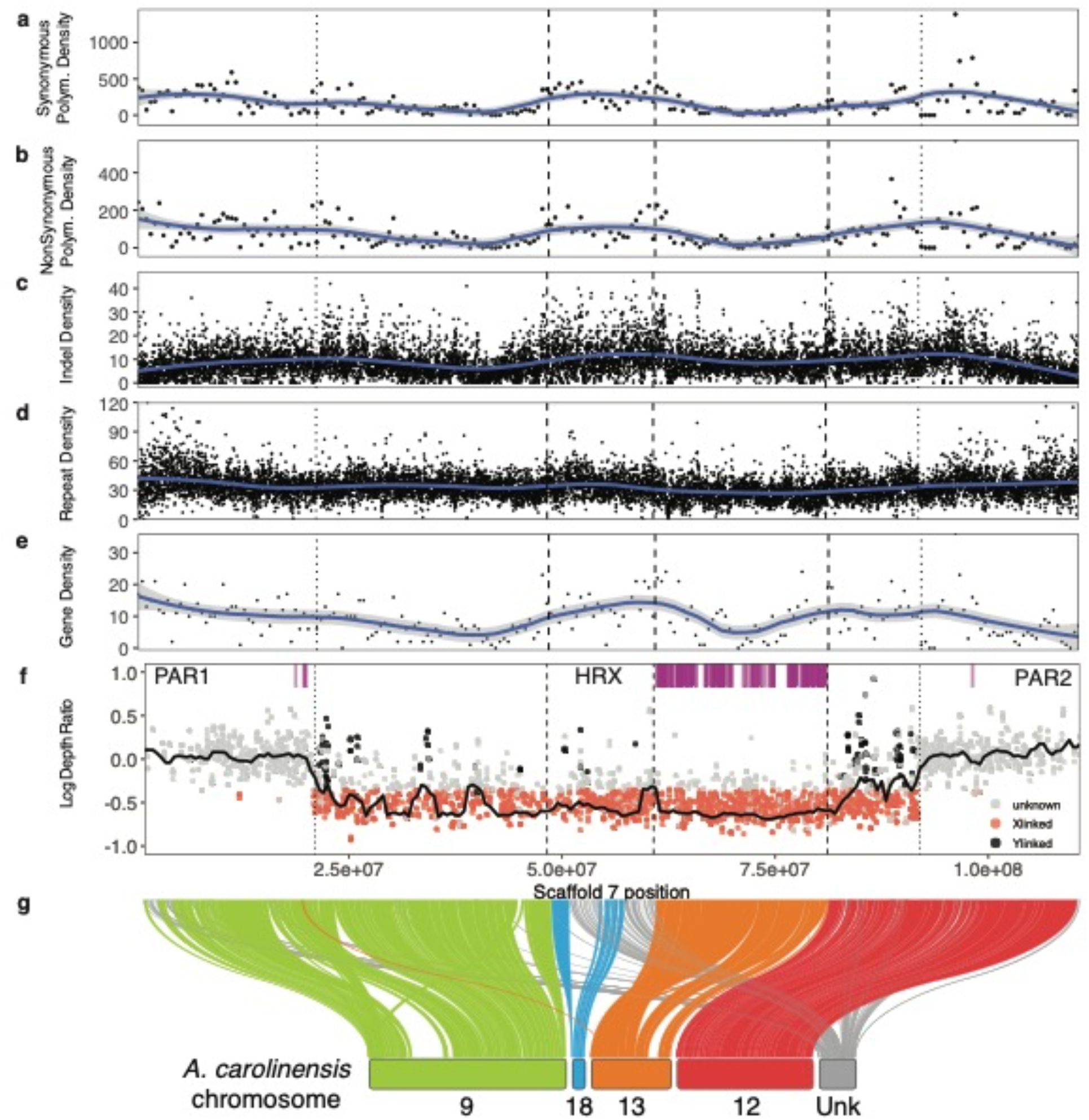
Identification and analysis of the X chromosome. **a-e)** sliding window plots of element density with LOESS smoothed lines (span=0.25) of **a**) synonymous SNPs per 500kb, **b**) non-synonymous SNPs per 500kb, **c**) indels per 10kb, **d**) Repetitive elements per 10kb, and **e**) Genes per 500kb. **f**) Scaffold 7 male/female depth ratio (log-transformed). The black horizontal line summarizes depth ratio using a sliding window analysis (2Mb windows, 500 kb step size). X-linked SNPs as those that are outliers for low sequencing depth ratio and show male heterozygosity equal to or lower than female heterozygosity. Y-linked SNPs correspond to significant sex GWA hits. Magenta ticks indicate the annotated location of *A. sagrei* homologs of X-linked genes in *A. carolinensis*. **g**) Syntenic relationship between *A. sagrei* scaffold 7 and *A. carolinensis* chromosomes. For all panels, dashed lines represent the boundaries between regions homologous to different A. carolinensis chromosomes and dotted lines mark the estimated boundaries of pseudoautosomal regions (PAR1 and PAR2) and the putative Hemizygous Region of the X (HRX).

Although our synteny data confirmed the identity of the ancestral chromosomes that fused to the ancient X to make up *A. sagrei*’s neo-sex chromosomes, our results do not support previous predictions of how these chromosomes are ordered within the *A. sagrei* neo-X chromosome. Our data suggest that chromosomes 18 and the ancient X are fused together at the center rather than at the extremes of scaffold 7 (Fig. 4g). In addition, only minor rearrangements are evident within formerly autosomal chromosomes, which suggests high levels of synteny within chromosomes despite their fusion to each other and the ancient X (Fig. 4).

We found that all 9 linkage groups that had been previously assigned to the ancient X aligned to a 20.35 Mb stretch near the center of scaffold 7 (Table S4). LGb, the first region in the *A. carolinensis* genome identified as X-linked, and GL343282.1 are the only linkage groups with more than one alignment to scaffold 7. LGb’s other alignment is relatively short (∼1.6 Mb) and is located in one of two hypothesized pseudoautosomal regions (see below); GL343282.1’s other hit, on the other hand, is also within the boundaries of the region homologous to the ancient X (see Table S5). In addition to partially corroborating Kichigin and colleagues^56^ hypothesis and predicting a new order for the ancestral autosomes along *A. sagrei*’s neo-sex chromosome system (including the ancient X), our syntenic alignment also identified an additional 142 linkage groups from the *A. carolinensis* assembly as being X-linked in *A. sagrei*’s scaffold 7 (Table S5).

Our results, therefore, corroborate the hypothesis that the XY system in *A. sagrei* is composed of neo-sex chromosomes that originated through the fusion of chromosomes homologous to chromosomes 9, 12, 18 and the X in the *A. carolinensis* karyotype^56^. Furthermore, the high contiguity of scaffold 7 led us to hypothesize a new arrangement of these formerly autosomal chromosomes in the *A. sagrei* neo-X chromosome.

### Sex Chromosome Identification

Previous studies have indicated that *A. sagrei* has a male heterogametic sex chromosome system^68,69,76^. The sex chromosomes of this species are thought to be represented either by microchromosomes^76,77^ or by macrochromosomes^62^. Our synteny-based analyses (above) suggest that scaffold 7 is the X chromosome in *A. sagrei*. To independently verify which of the chromosomes in the *A. sagrei* genome are sex-linked, we used double-digest restriction site associated (ddRAD) data for 50 males and 50 females drawn from 16 populations distributed across the native and introduced ranges of *A. sagrei* (Table S6). This method has previously been shown to perform well for sex chromosome identification in anoles^78^ and other taxa^79^.

After quality filtering, we retained an average of 2.3 M read pairs per sample, with no difference observed for males and females (*P* = 0.81; Wilcoxon rank sum test). The GWAS analysis performed using the final 120,967 filtered SNP set identified 204 markers distributed on scaffolds 1, 2, 3, 5, 6, and 7 as significantly associated with sex (Fig. S1a). Of these, the majority (i.e., 190 SNPs; 93.1%) were clustered on scaffold 7, where we also identified the strongest associations (Fig. S1a, b). Most (96%) significant GWAS hits showed an excess of heterozygosity in males relative to females (Fig. S1c, d), as expected if they are linked with the Y chromosome. Compared to the significant associations on scaffold 7, those occurring on scaffolds 1-6 showed an excess of sequencing coverage in males relative to females (Fig. S1c, d,e). Therefore, a reasonable interpretation is that these SNPs correspond to regions of the genome that have been duplicated between the autosomes and the Y chromosome.

Analysis of sequencing depth further supported our interpretation that scaffold 7 is the sex chromosome in *A. sagrei*. Specifically, scaffold 7 contains 89.6% of the genomic outliers with lower coverage in males compared to females (Fig. 4g). This result is consistent with heterogamety in *A sagrei*, and with X-linkage of sequencing depth outliers. X-linked SNPs are clustered along a 71 Mb region on scaffold 7, which also contains the Y-linked SNPs identified by GWAS (Fig. S1b).

Collectively, these results indicate the 71 Mb region on scaffold 7 corresponds to the putative Hemizygous Region of the X chromosome (HRX) in *A. sagrei*. The 21 Mb to the left of the HRX, and the 19 Mb to the right of the HRX mostly contain markers with even coverage between males and females. We infer that these correspond to recombining pseusoautosomal regions (PAR1 and PAR2; Fig. 4f). These pseudoautosomal regions (PARs) appear to have evolved *de novo* since the divergence of *A. sagrei* and *A. carolinensis* as virtually all the ancestral X chromosome (and therefore the ancestral PARs) lies outside of the PAR in *A. sagrei*. A similar – but far more ancient – event has been hypothesized to have occurred in eutherian mammals where one of the two PARs present in these species arose after the divergence of monotremes and placental mammals 80-130 million years ago^80^. Two recent studies place the divergence of *A. sagrei* and *A. carolinensis* at less than 50 million years ago^81,82^ suggesting that *A. sagrei* has evolved two new PARs in roughly half the time placental mammals evolved one.

As discussed above, we detected broad sequence homology between *A. sagrei* scaffold 7 and nine X-linked *A. carolinensis* scaffolds which together contain 272 gene models in the NCBI RefSeq^83^ *A. carolinensis* annotation (release 102). We found 227 orthologous gene models in our annotation of *A. sagrei*. The vast majority (224; 99%) of these appear on scaffold 7 in AnoSag2.1. Only three genes are annotated to occur on other scaffolds (*gal3st1, iscu*, and *iqcd* on scaffolds 2,6, and 9 respectively). Most of these genes occur exclusively on scaffold 7, however, 17 genes have paralogs occurring both on scaffold 7 and another scaffold. Of the 224 genes on scaffold 7, all of them have at least one copy within the region homologous to the *A. carolinensis* X, chromosome 13 (Fig 4f). Duplicate copies of two genes also occur elsewhere on scaffold 7. A single copy of *dnah10* is present in PAR1 and three copies of *cmklr1* occur in PAR1 and one copy in PAR2 (Table S7). In mammals and flies the duplication or movement of genes to regions outside HRX have been observed and have been hypothesized to be associated with either dosage compensation or male-specific function. However, we are unable to find support for either hypothesis for these two genes.

### X-autosome fusion

The Y chromosome of *A. sagrei* is roughly two-thirds the size of the X^62^. This reduction is likely via the process of Y chromosome degendration^55,84^. Under this process, formerly homologous regions in the Hemizygous Region of the X chromosome (HRX) of the X and Y diverge through mutational accumulation and deletions on the Y. The HRX is expected to evolve under different evolutionary pressures than those on autosomes or within pseudoautosomal regions on the sex chromosomes because, when they occur in males, these loci are effectively haploid. Recessive deleterious genetic variants such as indels, non-synonymous mutations, or repetitive element insertions are thus exposed to purifying natural selection in males and are therefore more likely to be purged from a population^85^. Similarly, the hemizygosity of the X chromosome may result in more efficient positive natural selection^86,87^. Since the *A. sagrei* neo-sex chromosomes are composed of ancient sex-linked sequences as well as more recently recruited former autosomes, we might expect variation in the density of variants among these regions, reflecting differences in the time they have been X-linked. Just such a phenomenon has been observed in the neo-X of *Drosophila miranda* where formerly autosomal portions of the X chromosome have reduced synonymous polymorphism due to repeated selective sweeps^88^. Indeed, our data suggest some gametologs on the X and Y have sufficiently diverged to allow detection of X- and Y-linked sequences in the HRX of *A. sagrei* (Fig 4f). However, the mapping of male-linked sequences to regions homologous to *A. carolinensis* chromosomes 9, 12, and 18 but not the X (chromosome 13) reveals, unsurprisingly, that X-Y divergence is more substantial on the portion of the X chromosome that has been sex-linked the longest. We also observed differences in the density of indels, repetitive elements, genes, and synonymous and nonsynonymous polymorphisms among the sub-compartments of the *A. sagrei* X chromosome; regions homologous to the ancient X have a lower density of each of these features than regions homologous to *A. carolinensis* autosomes (Figs 4a-e, S2). Future analyses, using population genetic data in contrast to the pooled sequencing performed here, would allow more detailed evaluation of the evolutionary dynamics at play on the *A. sagrei* X chromosome.

## Concluding Remarks

We report a new, high-quality genome assembly of the brown anole, *Anolis sagrei*. Our analyses of this genome have revealed new insights into the lineage-specific accumulation of repetitive elements and the complex evolution of anole sex chromosomes, including multiple bouts of autosome-sex chromosome fusion. The highly contiguous nature of our assembly and its substantial completeness presents a community resource that will enable future and on-going work in this emerging model organism. The assembly and accompanying annotation of genes and genetic variation we report here make possible a wide array of analyses such as genetic mapping of traits (Bock et al. *accepted*, Feiner et al. in review) and functional genetics. Finally, the assembly serves as a launching point for future work probing the genome of this diverse species, including the assembly of the Y chromosome and population-scale analysis of structural evolution.

## Methods

### Chosen Animal

A single female *Anolis sagrei ordinatus* was chosen for sequencing. This animal was collected from the Conception Island Bank in the Eastern Bahamas. Mitochondrial sequencing from across the range of the species had previously revealed this population to have the lowest levels of nucleotide polymorphism^89^ and was therefore best suited for *de novo* genome assembly. After humane euthanasia using Sodium Pentobarbital, we excised and flash froze muscle and liver tissue in liquid nitrogen. Flash frozen tissues were subsequently stored at -80°C. All animal work was performed under Harvard Institutional Animal Care and Use Committee Protocol 26-11. Research, collection, and export permissions were granted by the Bahamas Environment, Science and Technology Commission, the Bahamas Ministry of Agriculture and Marine Resources, and the Bahamas National Trust.

### Sequencing

High Molecular Weight DNA was extracted from muscle and liver tissues using a Qiagen genomic tip kit. Two whole genome shotgun sequencing libraries were prepared using a TruSeq v3 DNA PCR-free library preparation kit with a 450bp insert between pairs.

Two Chicago libraries and three Dovetail HiC libraries were prepared following previously published protocols^59,90^. For Chicago libraries, ∼500ng of DNA was reconstituted into chromatin *in vitro* and then fixed in formaldehyde. For HiC libraries chromatin was first fixed in place with formaldehyde in the nucleus and then extracted. The remaining steps for both protocols were identical. Fixed chromatin was digested with DpnII, creating 5’ overhangs which were filled with biotinylated nucleotides followed by ligation of free blunt ends. Crosslinks were then reversed, and the DNA purified from protein. Purified DNA was treated to remove biotin that was not internal to ligated fragments. The DNA was then sheared to an average fragment size of 350 bp and used to generate sequencing libraries using NEBNext Ultra enzymes and Illumina-compatible adapters. Biotin-containing fragments were isolated using streptavidin beads before PCR enrichment of each library.

The two whole genome shotgun (WGS) libraries were multiplexed and sequenced across two sequencing lanes. The two Chicago and three HiC libraries were multiplexed and sequenced across two additional lanes. All libraries were sequenced as paired end 150bp reads on the Illumina HiSeqX platform. A summary of the data generated from all sequencing approaches can be found in Table S8.

### de novo Assembly

We processed raw Illumina WGS reads using trimmomatic v.0.36^91^. We used ILLUMINACLIP to remove TruSeq3 v2 sequencing adapters. We then removed any nucleotides with quality scores less than 20 from the leading and trailing ends of each read. Next, reads were truncated from the ends if sliding windows of 13bp have an average quality below 20. Finally, we retained only reads longer than 23 nucleotides. For trimmed reads less than 23bp we removed both that read and its paired read. We retained 896 million read pairs after filtering. These reads were used as input for *de novo* assembly using meraculous v2.2.2.5^58^ with the following parameters (diploid mode - diploid nonredundant haplotigs, kmer size 73, minimum kmer frequency 8).

### Scaffolding

We used the initial *de novo* assembly, Chicago library reads, and Dovetail HiC library reads as input data for HiRise v2.1.6-072ca03871cc, a software pipeline designed specifically for using proximity ligation data to scaffold genome assemblies^59^. We performed an iterative process of scaffolding. First, Chicago library sequences were aligned to the *de novo* input assembly from meraculous using a modified SNAP read mapper (http://snap.cs.berkeley.edu). The mapped separation of Chicago read pairs within draft scaffolds were analyzed by HiRise to produce a likelihood model for genomic distance between read pairs, and the model was used to identify and break putative misjoins, to score prospective joins, and make joins above a threshold. After aligning and scaffolding using Chicago data, HiC library sequences were aligned and used for scaffolding following the same method above but with the Chicago-scaffolded assembly as input.

### AnoSag1.0

Using the Chicago-scaffolded assembly as input we used abyss-sealer v2.02^92^ with options “`-v - j32 -s100G -k96 -k80 -k64 -k48 -P 50 -o run20 -B5000`” to close 18.6% of gaps in the assembly, substituting 9 Mb of ambiguous sequence with determined bases and increased N50 by 1.2Mb. Gap-filled scaffolds were ordered and named sequentially according to descending size.

The total length of this assembly (1.6 Gb) was substantially smaller than the *Anolis carolinensis* assembly or cytological estimates of *A. sagrei* (∼1.9 Gb), so we performed an additional round of improvement to identify missing sequences. Using bwa-mem v0.17^93^, we mapped the Illumina WGS PE reads generated in the project to the v1.0 assembly and extracted all unmapped reads. Approximately 2.5% of all read pairs either did not map or only a single read mapped. We then performed a *de novo* assembly using ABySS v2.02^94^ using the unmapped paired and unpaired reads as input and a kmer size of 96, generating ∼29K contigs. BLASTN v2.7.1^95^ annotation of these contigs against the NCBI nr database revealed that about half had a highest match to saurian sequences (14,360; of which 10,975 mapped to an *Anolis* accession). We reserved all contigs mapping to saurians and discarded all other contigs to avoid contaminant and metagenomic sequences. These contigs were appended to the final sagrei assembly and are numbered in descending size. The version 1.0 assembly including both gap-filled scaffolds and newly assembly and filtered contigs was composed of 28,096 elements (scaffolds plus contigs) totaling 1.62Gb in length.

### AnoSag2.0

During quality control checks of the v1.0 assembly, we discovered an issue that led us to further refine the assembly. Specifically, while spot checking the annotation of deeply conserved developmental genes, we discovered that while our assembly placed exons in the same order as other vertebrate genomes the orientation of exons within genes varied substantially. This seems to be caused by the inability of scaffolding software to determine the orientation of some contigs while performing scaffolding using Chicago and HiC data^96^. To correct this issue, we generated additional long read Pacific Biosciences data, broke the assembly back into contigs, and re-scaffolded.

### High Molecular Weight DNA isolation and sequencing

High-molecular-weight genomic DNA was extracted from the muscle tissue of the same female *A. sagrei ordinatus* used for all previous sequencing. Frozen muscle tissue was homogenized with a pestle in freshly made lysis buffer (0.1 mM Tris, 1% Polyvinylpyrrolidone 40, 1% Sodium metabisulfite, 500 mM NaCl, 50 mM EDTA, 1.25% SDS, pH 8.0) and incubated with proteinase K at 55°C for 50 minutes prior to RNase A treatment at room temperature for 10 minutes^97^. Next, one-third volume of 5M potassium acetate was added, and the solution was incubated at 4°C for 5 minutes and pelleted by centrifugation. The DNA in the supernatant was bound to SPRSelect beads (Beckman Coulter Life Sciences), washed and eluted in elution buffer (10 mM Tris, pH 8.0) according to the manufacturer’s instructions. Throughout the extraction process, the solutions were manipulated gently to minimize shearing of DNA.

### PacBio Sequencing

A SMRTbell library was constructed using the SMRTbell Template Prep Kit 1.0 (Pacific Biosciences) and sequenced on the Sequel I platform using Sequel Sequencing Kit 2.1 (Pacific Biosciences, Sequel SMRT Cell 1M v2). Two sequencing runs generated a total of 1,257,251 reads, with an average size of 18 kb.

### PacBio Contig Exension and Bridging

The Illumina short read data generated for the initial *de novo* assembly (see above) were used to correct errors in PacBio long reads using Proovread v2.14.1^98^. As a trade-off between run time and accuracy, 40x short read coverage was used during error-correction. The resulting untrimmed error-corrected PacBio reads were subjected to additional hybrid error correction with FMLRC^99^ before being used to extend and bridge the original contigs from the AnoSag1.0 assembly. The AnoSag1.0 genome assembly was reverted to contigs by breaking scaffolds at any gap of 100bp or more. The resulting contigs were extended and bridged using error-corrected PacBio reads using SSPACE-LongRead scaffolder v1.1^100^. Redundant contigs were removed using fasta2homozygous.py, a python script from Redundans v0.14a^101^. BUSCO assessments using the vertebrata dataset (vertebrata_odb9, containing 2586 highly conserved single-copy core vertebrate genes) were performed before and after the removal of redundant contigs to ensure that removing redundant contigs did not change the completeness of the genome assembly. Gaps in contigs were closed by LR Gapcloser^102^. Contigs were then re-scaffolded through two iterations of the HiRise pipeline using previously prepared Chicago and Hi-C libraries as described above to generate the version 2 *Anolis sagrei* genome assembly – AnoSag2.0. All programs were run using the recommended default settings.

### AnoSag2.1

We generated a link density histogram paired reads from our HiC library using Juicer v1.6^61^. Visualizing these data in Juicebox v1.11.08^103^ revealed the second largest scaffold was in fact an intercalated fusion of two large scaffolds (Fig. S3). Through inspection of HiC link data as well as read mapping of Illumina short-read, RNA-Seq, and PacBio data, we identified three breakpoints on the AnoSag2.0 scaffold_2 (positions 1-206,031,901; 206,031,902-209,142,770; 209,142,7701-210,003,944; and 210,003,945-342,856,123) resulting in 4 fragments. We split the scaffolds at these locations and rejoined the first fragment to the third, and the second fragment to the fourth according to evidence from HiC link mapping. This resulted in the formation of two new scaffolds – the fifth and sixth largest in the new assembly. We sorted and renamed scaffolds by size to create the final AnoSag2.1 assembly reported here. We are making earlier assemblies (AnoSag1.0 and AnoSag2.0) publicly available because earlier research has been performed and published using those preliminary assemblies.

### Mitochondrial Genome assembly

The mitochondrial genome was absent from the AnoSag2.1 assembly. To assemble the mitochondrial genome, we first subsampled 1 million trimmomatic filtered and trimmed Illumina read pairs. These reads were used as input for a circular *de novo* assembly in Geneious v11.1.5 (https://www.geneious.com). Separately, we extracted the largest subread (17.2 kb) from our error-corrected PacBio dataset. These two sequences were identical at the nucleotide level where they overlapped, but each contained regions absent in the other. To complete the mtGenome assembly we created a consensus of these two sequences and then aligned both PacBio and Illumina reads to that consensus to confirm reads from both platforms aligned to all regions. We annotated the mitochondrial genome assembly using the MITOS webserver^104^.

### Chromosome Size analysis

Using a recently published, high-resolution *Anolis sagrei* karyotype (Figure 1c from Giovannotti and colleages^62^) we measured the size of each chromosome as it appeared in that figure. For each chromosome, we calculated the fraction of the total karyotype occupied by that chromosome for an XX individual and multiplied that fraction by the total size of the AnoSag2.1 assembly to generate an estimate of nucleotide content. These estimates were then compared against the number of nucleotides in each size sorted AnoSag2.1 scaffold (Table S9). We calculated the correlation between scaffold size and estimated chromosome size via linear regression using the lm function in R v3.6^105^.

### Repetitive Element Content

To estimate the repetitive landscape of the *Anolis sagrei* genome, we modeled repeats *de novo* on the assembly using RepeatModeler v1.08^106^ and annotated the repeat consensus sequences using RepeatMasker v4.0.7^107^. To understand the age distribution of transposable elements in each genome, we used the divergence of an insert from its family consensus as a proxy for its age. We generated alignments for each repeat family and calculated the Kimura-2 parameter divergence from consensus (correcting for CpG sites) using the calcDivergenceFromAlign.pl RepeatMasker tool. We compared the repetitive profiles of *A. sagrei* and *A. carolinensis* through a parallel analysis, running RepeatModeler and RepeatMasker with the AnoCar2.0 assembly^19^.

### Gene Model Annotation

For gene structure annotation of AnoSag2.1, we ran Braker v2.0.5^67^ using RNA-Seq data and amino acid sequences of closely related species. In brief, we used RepeatModeler v1.0.11^106^ to construct an *Anolis sagrei* repeat library, which was subsequently used by RepeatMasker v1.0.11^107^ to mask repeats in the genome. We used protein sequences of *A. carolinensis* and *A. punctatus* obtained from NCBI RefSeq to query our reference sequence for homologous proteins. Composite RNA-seq data were prepared by combining eight paired-end RNA-Seq libraries consisting of two libraries from a forelimb and a hindlimb at embryonic stage 7^108^, three libraries from brain, liver, and skin tissue of an adult female^109^ (SRA accession number: DRA004457), and three libraries from central, nasal, and temporal regions of eye retina at embryonic stage 16.5. These RNA-seq reads were aligned to AnoSag2.0 using TopHat v2.1.1^110^ with the option -- b2-very-sensitive^110^. The Braker gene prediction pipeline was run with the options “`-- softmasking --prg=gth --gth2traingenes`”.

CD-HIT v4.6.8^111,112^ was used with default parameters to remove redundant gene models from Braker’s output. Using the protein sequences of non-redundant gene models from CD-HIT as a query, BLASTP v2.7.1^95^ searches were performed against the non-redundant RefSeq protein database. Gene models with unique protein matches and e-value less than 1e-3 were kept. When more than one gene model had blast hits with the same protein, the gene model with the best score was kept. In addition, we retained gene models that lacked a blast hit if they either 1) contained 3 or more exons or 2) had more than 50 RNA-seq reads per 15 million mapped reads and did not overlap with those from the non-redundant CD-HIT gene models already retained. Gene models from these processes were combined to generate a final non-redundant gene set. Approximately 15% of final gene models were spot checked and manually edited by cross-referencing Braker gene model annotations with aligned RNA-seq data.

### SNP/Indels Genotyping

We performed sequence variant calling with composite shotgun data by combining 75bp single-end Illumina reads from 5 genomic libraries originally generated as control data for *A. sagrei* ChIP-seq experiments. Each genomic library was created from a pool of embryos produced by a colony of wild-caught *A. sagrei* from Orlando, FL. The library details are as follows: library 1, 57 embryos, 46.6 million reads; library 2, 59 embryos, 42.1 million reads; library 3, 91 embryos, 16.6 million reads; library 4, 97 embryos, 104 million reads; library 5, 70 embryos, 25.3 million reads (need to submit to GEO). Likewise, composite RNA-Seq data were generated by combining data from 27 RNA-Seq libraries from embryonic (forelimbs, hindlimbs, and retina) and adult tissues (brain, skin, and liver). Embryonic limb (GEO accession GSE128151) and adult tissue RNA-seq data (DDBJ Sequence Read Archive accession DRA004457) were previously published^108,109^. Anole eye RNA-seq datasets were generated from tissues from Sanger Stage 16.5 embryos laid from wild-caught *A. sagrei* parents from Orlando, FL. Tissues from the nasal, central, and temporal posterior regions of the eye were collected, and samples from 3 embryos (of mixed sex) were pooled together. Total RNA was isolated using the mirVana RNA Isolation Kit (ThermoFisher Scientific). Libraries were constructed with TruSeq Stranded mRNA Sample Prep Kit for Illumina and sequenced on the Illumina NextSeq 500 platform. Embryonic retina RNA-seq data were submitted to GEO (GEO accession GSE184570). The composite sequence data were aligned to the *A. sagrei* genome (AnoSag2.1) using BWA-mem v0.7.15^93^ with the default parameters. The resulting alignment files (SAM format) were merged and converted into sorted BAM file using SAMtools v1.6^113^ before duplicates were removed using Picard v2.16.0 (Broad Institute, 2019) MarkDuplicates with the options “`MAX_FILE_HANDLES_FOR_READ_ENDS_MAP=1000 REMOVE_DUPLICATES=true` `ASSUME_SORTED=true VALIDATION_STRINGENCY=LENIENT`”. SAMtools mpileup was used to generate genotype likelihoods at each genomic position with coverage from the deduplicated BAM file. We used BCFtools v1.9^114^ with the options “`--keep-alts --multiallelic-caller --variants-only`” to call and filter sequence variants. We further filtered single nucleotide variants using VCFTools v0.1.15^115^ to have a minimum quality score of 25 and a minimum depth of 5 reads (commands “`--minQ 25 --minDP 5`”). Summaries of features per genomic window (indels, SNPs, genes, repetitive elements) were calculated using VCFTools and BEDTools v2.26^116^. The impact of single nucleotide variants was assessed using SNPeff v5.0^117^ using default settings.

### Analysis of X chromosome synteny

We used SatsumaSynteny v2.0^118^ to align scaffold 7 in the *A. sagrei* assembly to the *A. carolinensis* assembly version 2.0. Next, we used custom awk scripts to modify SatsumaSynteny’s default output to a bed format and used the Circos^119^ function bundlelinks to merge adjacent links together. We used bundlelinks’ ‘strict’ flag, keeping only bundles that were within 1Mbp of each other and that were at least 1kbp in length. To make the linear synteny plot between *A. sagrei* scaffold 7 and the scaffolds in the *A. carolinensis* assembly, we used the R package RIdeogram^120^ in R v3.6^105^. For plotting purposes, we labeled scaffolds that aligned to scaffold 7 as belonging to either chromosome 9, 12, 18 or the ancient X in the *A. carolinensis* assembly using information from previous flow sorting and dosage compensation studies in *A. carolinensis*^*11,62,74,75,121*^.

We assessed the synteny degree between the *A. sagrei* and *A. carolinensis* genomes using Satsuma v3.1.0^118^, an alignment software devised to deal with large queries and references. Satsuma works as follows. First, it breaks the query and the reference sequences into 4096bp chunks, by default, that overlap in one-quarter of their size. Then, it translates As, Cs, Ts and Gs into numeric signals that go through cross-correlation. Cross-correlation is calculated as a function of the displacement of one sequence’s signal relative to the other. It measures the similarity among two analog signals, the higher the cross-correlation value, the more bases match across the overlap and the stronger the signal. Next, Satsuma fine-tunes the alignment by keeping sequences that are at least 28 base-pairs long and have 45% matching. Then, Satsuma calculates an alignment probability model based on the aligned sequences length, identity, GC content and length, keeping alignments that have probability lower than 10^−5^ of being random noise. After Satsuma identifies it proceeds with dynamic programming to merge overlapping blocks into alignments with gaps. To reduce computational time, Satsuma implements a ‘paper- and-pencil game battleship’ approach, in which it queries the vicinity of the alignment for more hits.

### Sex chromosome identification

Libraries were made following the standard ddRAD protocol^122^. Briefly, we used the *SphI* HF and *EcoRI* HF restriction endonucleases (New England Biolabs) to digest genomic DNA. After size selection, we retained fragments of 500–660 bp. Libraries were pair-end sequenced (150 bp read length) on an Illumina HiSeq 4000 (Illumina, San Diego, CA, USA). We included ddRAD data for 50 males and 50 females, sequenced as part of a larger related project. These were obtained from 16 populations distributed across the native and introduced ranges of *A. sagrei* (Table S6). In selecting samples, we aimed for a balanced representation of both sexes for most populations.

Sequencing files were de-multiplexed using *ipyrad* v0.7.15^123^. We removed low-quality bases and Illumina adapters using Trimmomatic v0.36^91^. Cleaned reads were used for SNP calling within the dDocent v2.2.20 pipeline^124^. In dDocent, reads were aligned to the *A. sagrei* assembly using BWA v0.7.16a-r1181^125^ at default parameters. We then performed joint variant calling using the 100 *A. sagrei* genotypes along with 925 other conspecific genotypes sequenced as part of related projects, in Freebayes v1.0.2^126^. The genotype calls for the 100 samples used here were filtered using vcflib (https://github.com/vcflib/vcflib). We kept only biallelic SNPs with MAPQ scores > 20. For the remaining markers, we coded genotypes that were supported by fewer than four reads as missing data. We subsequently kept only SNPs with data in at least 70% of samples, and those with minor allele frequency larger than 5%.

To identify Y-linked genomic regions, we performed a genome-wide association study (GWAS) in PLINK v1.0.7^127^. Because *A. sagrei* is known to have a male heterogametic sex chromosome system^62,68,76^, SNPs associated with one sex should be in close linkage with the Y chromosome. Specifically, we expect these SNPs to represent differences that occur between X and Y gametologs. We coded sex as a binary case/control variable. Prior to the GWAS analysis, we imputed any remaining missing data in the filtered SNP set using BEAGLE v5.0 at default parameters^128^. For association testing, we used Fisher’s exact test and set the genome-wide significance threshold using the Bonferroni correction for multiple comparisons (0.05/total number of tested markers). Further confirmation of GWAS results was obtained by calculating the difference in heterozygosity between males and females (i.e. relative male heterozygosity), at each SNP. This is because in a male-heterogametic system we expect SNPs occurring between gametologs to show an excess of heterozygosity.

To identify X-specific genomic regions, we used the ratio of sequencing coverage between males and females. In species with heteromorphic sex chromosomes such as *A. sagrei*, this metric should be effective in distinguishing between the autosomes and the X chromosome^79^. A ratio close to 1 is expected for autosomes, while a ratio of 0.5 is expected for the X chromosome, due to hemizygosity of males. For both sequencing depth ratio and relative male heterozygosity, we identified upper and lower thresholds for categorizing SNPs as genomic outliers using the interquartile range (IQR; upper/lower quartile +/- 1.5 IQR). We then defined X-linked SNPs as those that are outliers for low sequencing depth ratio and show male heterozygosity equal to or lower than female heterozygosity. Y-linked SNPs were defined as those with significant sex GWA hits. Lastly, we used these SNP categories to identify the approximate boundaries of the PARs, by tallying the percent of sex-linked SNPs in 1Mb windows along scaffold 7 (Table S10).

## Acknowledgements

Research, collection, and export permissions were granted by the Bahamas Environment, Science and Technology Commission, Bahamas National Trust, and the Bahamas Ministry of Agriculture and Marine Resources. Thank you to Alexis Harrison, Sofia Prado-Irwin, Shea Lambert, Dan Scantlebury, and Shannan Yates for their assistance in the field. We are grateful to Cory Hahn, Matthew Gage, Analisa Shields-Estrada, Jeff Breeze, and the postdoctoral fellows, graduate, and undergraduate students of the Losos lab for assisting with animal care. Thank you to Tonia Schwartz for providing advanced access to the *Sceloporus undulatus* genome for completeness and contiguity comparisons. All animal care procedures were approved by Harvard Institutional Animal Care and Use Committee Protocol 26-11. This work was supported by the National Science Foundation under grants DEB-1927194 to JBL and AJG, IOS-1827647 to DBM, and DEB-1927156 to DBM. Additionally, DGB was supported by a Natural Sciences and Engineering Research Council of Canada (NSERC) Postdoctoral Fellowship, and a Banting Postdoctoral Fellowship. This project was also made possible, in part, through the support of a grant from the John Templeton Foundation and Human Frontiers grant RGP0030/2020 to JBL and DBM. The opinions expressed in this publication are those of the authors and do not necessarily reflect the views of the John Templeton Foundation.

## References

1 Miga, K. H. et al. Telomere-to-telomere assembly of a complete human X chromosome. Nature 585, 79–84 (2020).

2 Chang, C.-H. et al. Islands of retroelements are major components of Drosophila centromeres. PLOS Biology 17, e3000241, doi:10.1371/journal.pbio.3000241 (2019).

3 Rhoads, A. & Au, K. F. PacBio sequencing and its applications. Genomics, proteomics & bioinformatics 13, 278–289 (2015).

4 Schield, D. R. et al. The origins and evolution of chromosomes, dosage compensation, and mechanisms underlying venom regulation in snakes. Genome Research 29, 590–601, doi:10.1101/gr.240952.118 (2019).

5 Pasquesi, G. I. M. et al. Squamate reptiles challenge paradigms of genomic repeat element evolution set by birds and mammals. Nature Communications 9, 2774, doi:10.1038/s41467-018-05279-1 (2018).

6 Feiner, N. Evolutionary lability in Hox cluster structure and gene expression in Anolis lizards. Evolution Letters 3, 474–484, doi:10.1002/evl3.131 (2019).

7 Hodson, C. N., Jaron, K. S., Gerbi, S. & Ross, L. Evolution of gene-rich germline restricted chromosomes in black-winged fungus gnats through introgression (Diptera: Sciaridae). bioRxiv, 2021.2002.2008.430288, doi:10.1101/2021.02.08.430288 (2021).

8 Fan, H. et al. Chromosome-level genome assembly for giant panda provides novel insights into Carnivora chromosome evolution. Genome biology 20, 1–12 (2019).

9 Mahajan, S., Wei, K. H.-C., Nalley, M. J., Gibilisco, L. & Bachtrog, D. De novo assembly of a young Drosophila Y chromosome using single-molecule sequencing and chromatin conformation capture. PLoS biology 16, e2006348 (2018).

10 Rovatsos, M. & Kratochvil, L. Evolution of dosage compensation does not depend on genomic background. Mol Ecol 30, 1836–1845, doi:10.1111/mec.15853 (2021).

11 Rupp, S. M. et al. Evolution of Dosage Compensation in Anolis carolinensis, a Reptile with XX/XY Chromosomal Sex Determination. Genome Biol Evol 9, 231–240, doi:10.1093/gbe/evw263 (2017).

12 Rhie, A. et al. Towards complete and error-free genome assemblies of all vertebrate species. Nature 592, 737–746, doi:10.1038/s41586-021-03451-0 (2021).

13 Koepfli, K.-P., Paten, B., Genome, K. C. o. S. & O’Brien, S. J. The Genome 10K Project: a way forward. Annual review of animal biosciences 3, 57–111, doi:10.1146/annurev-animal-090414-014900 (2015).

14 Twyford, A. D. The road to 10,000 plant genomes. Nature Plants 4, 312–313, doi:10.1038/s41477-018-0165-2 (2018).

15 Tang, H. et al. Synteny and Collinearity in Plant Genomes. Science 320, 486–488, doi:10.1126/science.1153917 (2008).

16 Flicek, P., Keibler, E., Hu, P., Korf, I. & Brent, M. R. Leveraging the Mouse Genome for Gene Prediction in Human: From Whole-Genome Shotgun Reads to a Global Synteny Map. Genome Research 13, 46–54, doi:10.1101/gr.830003 (2003).

17 Montgomery, T. H. A study of the chromosomes of the germ cells of metazoa. (1901).

18 Losos, J. B. Lizards in an Evolutionary Tree: Ecology and Adaptive Radiation of Anoles. (University of California Press, 2009).

19 Alföldi, J. et al. The genome of the green anole lizard and a comparative analysis with birds and mammals. Nature 477, 587–591, doi:10.1038/nature10390 (2011).

20 Williams, E. E. The ecology of colonization as seen in the zoogeography of anoline lizards on small islands. The Quarterly Review of Biology 44, 345–389 (1969).

21 Henderson & Powell. Amphians and Reptiles of the West Indies. (2009).

22 Reynolds, R. G. et al. Phylogeographic and phenotypic outcomes of brown anole colonization across the Caribbean provide insight into the beginning stages of an adaptive radiation. Journal of Evolutionary Biology 33, 468–494, doi:10.1111/jeb.13581 (2020).

23 van Wagensveld, T. P. & Questel, K. Cuban Brown Anole (Anolis sagrei) established on Anguilla. Reptiles & Amphibians 25, 162–163 (2018).

24 Powell, R., Henderson, R. W. & Farmer, M. C. Introduced amphibians and reptiles in the greater Caribbean: Patterns and conservation implications. … of Caribbean island …, 63–144, doi:10.1163/ej.9789004183957.i-228.38 (2011).

25 Fisher, S. R., Del Pinto, L. A. & Fisher, R. N. Establishment of brown anoles (Anolis sagrei) across a southern California county and potential interactions with a native lizard species. PeerJ 8, e8937 (2020).

26 Batista, A., Ponce, M., Garcés, O., Lassiter, E. & Miranda, M. Silent pirates: Anolis sagrei Duméril & Bibron, 1837 (Squamata, Dactyloidae) taking over Panama City, Panama. Check List 15, 455 (2019).

27 Amador, L., Ayala-Varela, F., Nárvaez, A. E., Cruz, K. & Torres-Carvajal, O. First record of the invasive Brown Anole, Anolis sagrei Duméril & Bibron, 1837 (Squamata: Iguanidae: Dactyloinae), in South America. Check List 13, 2083 (2017).

28 Stroud, J. T., Giery, S. T. & Outerbridge, M. E. Establishment of Anolis sagrei on Bermuda represents a novel ecological threat to Critically Endangered Bermuda skinks (Plestiodon longirostris). Biological Invasions, 1–9, doi:10.1007/s10530-017-1389-1 (2017).

29 Stroud, J. T., Richardson, A. J., Sim, J., Airnes, A. & Stritch, J. M. Onward to the Mid-Atlantic: First records of Cuban brown anoles (Anolis sagrei) on Ascension Island. Reptiles & Amphibians 25, 220–222 (2018).

30 Goldberg, S. R., Kraus, F. & Bursey, C. R. Reproduction in an introduced population of the brown anole, Anolis sagrei, from O’ahu, Hawai’i. Pacific Science 56, 163–168, doi:10.1353/psc.2002.0014 (2002).

31 Sanger, T. J., Pm, H., Johnson, M. A., Diani, J. & Losos, J. B. Laboratory Protocols for Husbandry and Embryo Collection of Anolis Lizards. Herpetological Review 39, 58–63, doi:papers3://publication/uuid/2E4CD7E0-5C43-4465-AE59-0C32CA979A05 (2008).

32 De Meyer, J. et al. in th Anole Newsletter (eds James T. Stroud, Anthony J. Geneva, & Jonathan B. Losos) 1–26 (2019).

33 Losos, J. B. Integrative Approaches to Evolutionary Ecology: Anolis Lizards as Model Systems. Annual Review Of Ecology And Systematics 25, 467–493, doi:10.1146/annurev.es.25.110194.002343 (1994).

34 Losos, J. B. & Pringle, R. M. Competition, predation and natural selection in island lizards. Nature 475, E1–E2, doi:papers3://publication/doi/10.1038/nature10140 (2011).

35 Pringle, R. M. et al. Predator-induced collapse of niche structure and species coexistence. Nature 570, 58–64, doi:10.1038/s41586-019-1264-6 (2019).

36 Schoener, T. W. & Schoener, A. Ecological and demographic correlates of injury rates in some Bahamian Anolis lizards. Copeia 1980, 839, doi:papers3://publication/doi/10.2307/1444463 (1980).

37 Lapiedra, O., Schoener, T. W., Leal, M., Losos, J. B. & Kolbe, J. J. Predator-driven natural selection on risk-taking behavior in anole lizards. Science 360, 1017–1020, doi:10.1126/science.aap9289 (2018).

38 Lapiedra, O., Chejanovski, Z. & Kolbe, J. J. Urbanization and biological invasion shape animal personalities. Glob Chang Biol 23, 592–603, doi:10.1111/gcb.13395 (2017).

39 Sanger, T. J., Losos, J. B. & Gibson-Brown, J. J. A developmental staging series for the lizard genus Anolis: a new system for the integration of evolution, development, and ecology. Journal of Morphology 269, 129–137, doi:papers3://publication/doi/10.1002/jmor.10563 (2008).

40 Tiatragul, S., Kurniawan, A., Kolbe, J. J. & Warner, D. A. Embryos of non-native anoles are robust to urban thermal environments. J Therm Biol 65, 119–124, doi:10.1016/j.jtherbio.2017.02.021 (2017).

41 Warner, D. & Moody, M. Egg environments have large effects on embryonic development, but have minimal consequences for hatchling phenotypes in an invasive lizard. … Journal of the … 105, 25–41, doi:10.1111/j.1095-8312.2011.01778.x (2012).

42 Warner, D. & Moody, M. Is water uptake by reptilian eggs regulated by physiological processes of embryos or a passive hydraulic response to developmental environments? Comparative Biochemistry and … 160, 421–425, doi:10.1016/j.cbpa.2011.07.013 (2011).

43 Park, S., Infante, C. R., Rivera-Davila, L. C. & Menke, D. B. Conserved regulation of hoxc11 by pitx1 in Anolis lizards. J Exp Zool B Mol Dev Evol 322, 156–165, doi:10.1002/jez.b.22554 (2014).

44 D’Agostino, E. R. R., Donihue, C. M., Losos, J. B. & Geneva, A. J. ASSESSING THE EFFECTS OF GENETIC DIVERGENCE AND MORPHOLOGY ON ANOLIS LIZARD MATING. BioOne 568, 1–15, doi:papers3://publication/doi/10.3099/0006-9698-568.1.1.short (2020).

45 Driessens, T. et al. Climate-related environmental variation in a visual signalling device: the male and female dewlap in Anolis sagreilizards. Journal of Evolutionary Biology 30, 1846–1861, doi:10.1111/jeb.13144 (2017).

46 Tokarz, R. R., McMann, S., Smith, L. C. & John-Alder, H. Effects of testosterone treatment and season on the frequency of dewlap extensions during male-male interactions in the lizard Anolis sagrei. Horm Behav 41, 70–79, doi:10.1006/hbeh.2001.1739 (2002).

47 Baeckens, S., Driessens, T. & Van Damme, R. The brown anole dewlap revisited: do predation pressure, sexual selection, and species recognition shape among-population signal diversity? PeerJ 6, e4722, doi:10.7717/peerj.4722 (2018).

48 Kolbe, J., Leal, M. & Schoener, T. Founder Effects Persist Despite Adaptive Differentiation: A Field Experiment with Lizards. Science 335, 1086–1089, doi:papers3://publication/doi/10.1126/science.1209566 (2012).

49 Kolbe, J. J., Larson, A., Losos, J. B. & de Queiroz, K. Admixture determines genetic diversity and population differentiation in the biological invasion of a lizard species. Biology letters 4, 434–437, doi:papers3://publication/doi/10.1098/rsbl.2008.0205 (2008).

50 Kolbe, J. J. et al. Genetic variation increases during biological invasion by a Cuban lizard. Nature 431, 177–181, doi:papers3://publication/doi/10.1038/nature02807 (2004).

51 Bock, D. et al. Changes in selection pressure can facilitate hybridization during biological invasion. PNAS (in review).

52 Losos, J. B., Warheitt, K. I. & Schoener, T. W. Adaptive differentiation following experimental island colonization in Anolis lizards. Nature 387, 70–73, doi:10.1038/387070a0 (1997).

53 Kolbe, J. J., Leal, M., Schoener, T. W., Spiller, D. A. & Losos, J. B. Founder effects persist despite adaptive differentiation: a field experiment with lizards. Science 335, 1086–1089, doi:10.1126/science.1209566 (2012).

54 Logan, M. L., Cox, R. M. & Calsbeek, R. Natural selection on thermal performance in a novel thermal environment. Proc Natl Acad Sci U S A 111, 14165–14169, doi:10.1073/pnas.1404885111 (2014).

55 Lisachov, A. P. et al. Genetic Content of the Neo-Sex Chromosomes in Ctenonotus and Norops (Squamata, Dactyloidae) and Degeneration of the Y Chromosome as Revealed by High-Throughput Sequencing of Individual Chromosomes. Cytogenetic and Genome Research 157, 115–122, doi:papers3://publication/doi/10.1159/000497091 (2019).

56 Kichigin, I. G. et al. Evolutionary dynamics of Anolis sex chromosomes revealed by sequencing of flow sorting-derived microchromosome-specific DNA. Molecular Genetics and Genomics 291, 1955–1966, doi:10.1007/s00438-016-1230-z (2016).

57 Rasys, A. M. et al. CRISPR-Cas9 Gene Editing in Lizards through Microinjection of Unfertilized Oocytes. Cell Rep 28, 2288–2292 e2283, doi:10.1016/j.celrep.2019.07.089 (2019).

58 Chapman, J. A. et al. Meraculous: de novo genome assembly with short paired-end reads. PLoS One 6, e23501, doi:10.1371/journal.pone.0023501 (2011).

59 Putnam, N. H. et al. Chromosome-scale shotgun assembly using an in vitro method for long-range linkage. Genome Research 26, 342–350, doi:papers3://publication/doi/10.1101/gr.193474.115 (2016).

60 De Smet, W. The nuclear Feulgen-DNA content of the vertebrates (especially reptiles), as measured by fluorescence cytophotometry, with notes on the cell and chromosome size. Acta Zoologica et Pathologica Antverpiensia (Belgium) 76, 119–157 (1981).

61 Durand, N. C. et al. Juicer Provides a One-Click System for Analyzing Loop-Resolution Hi-C Experiments. Cell Syst 3, 95–98, doi:10.1016/j.cels.2016.07.002 (2016).

62 Giovannotti, M. et al. New insights into sex chromosome evolution in anole lizards (Reptilia, Dactyloidae). Chromosoma 126, 245–260, doi:10.1007/s00412-016-0585-6 (2017).

63 Andrade, P. et al. Regulatory changes in pterin and carotenoid genes underlie balanced color polymorphisms in the wall lizard. Proceedings of the National Academy of Sciences 116, 5633–5642, doi:10.1073/pnas.1820320116 (2019).

64 Westfall, A. K. et al. A chromosome-level genome assembly for the Eastern Fence Lizard (Sceloporus undulatus), a reptile model for physiological and evolutionary ecology. 213, 1–43, doi:10.1101/2020.06.06.138248 (2020).

65 Gemmell, N. J. et al. The tuatara genome reveals ancient features of amniote evolution. Nature, 1–26, doi:10.1038/s41586-020-2561-9 (2020).

66 Roscito, J. G. et al. The genome of the tegu lizard Salvator merianae: combining Illumina, PacBio, and optical mapping data to generate a highly contiguous assembly. Gigascience 7, 587–513, doi:10.1093/gigascience/giy141 (2018).

67 Hoff, K. J., Lomsadze, A., Borodovsky, M. & Stanke, M. Whole-Genome Annotation with BRAKER. Methods Mol Biol 1962, 65–95, doi:10.1007/978-1-4939-9173-0_5 (2019).

68 Gamble, T., Geneva, A. J., Glor, R. E. & Zarkower, D. Anolis sex chromosomes are derived from a single ancestral pair. Evolution 68, 1027–1041, doi:10.1111/evo.12328 (2014).

69 Rovatsos, M., Altmanová, M., Pokorná, M. & Kratochvíl, L. Conserved sex chromosomes across adaptively radiated Anolis lizards. Evolution 68, 2079–2085 (2014).

70 Bachtrog, D. A dynamic view of sex chromosome evolution. Current Opinion in Genetics & Development 16, 578–585, doi:10.1016/j.gde.2006.10.007 (2006).

71 Gu, L. & Walters, J. R. Evolution of Sex Chromosome Dosage Compensation in Animals: A Beautiful Theory, Undermined by Facts and Bedeviled by Details. Genome Biology and Evolution 9, 2461–2476, doi:10.1093/gbe/evx154 (2017).

72 Altmanová, M. et al. All iguana families with the exception of basilisks share sex chromosomes. Zoology (Jena) 126, 98–102, doi:10.1016/j.zool.2017.11.007 (2018).

73 Giovannotti, M. et al. New insights into sex chromosome evolution in anole lizards (Reptilia, Dactyloidae). Chromosoma, 1–16, doi:10.1007/s00412-016-0585-6 (2016).

74 Marin, R. et al. Convergent origination of a Drosophila-like dosage compensation mechanism in a reptile lineage. Genome Research 27, 1974–1987, doi:10.1101/gr.223727.117 (2017).

75 Rovatsos, M., Altmanová, M., Pokorná, M. J. & Kratochvíl, L. Novel X-linked genes revealed by quantitative polymerase chain reaction in the green anole, Anolis carolinensis. G3 (Bethesda) 4, 2107–2113, doi:10.1534/g3.114.014084 (2014).

76 De Smet, W. Description of the orcein stained karyotypes of 27 lizard species (Lacertilia, Reptilia) belonging to the families Iguanidae, Agamidae, Chameleontidae and Gekkonidae (Ascalabota). Acta Zool. Pathol. Antverpiensia 76, 35–72 (1981).

77 Gorman, G. C. & Atkins, L. Chromosomal heteromorphism in some male lizards of the genus Anolis. American Naturalist 100, 579–583, doi:papers3://publication/uuid/A515523A-C428-4A17-B24A-16297091656C (1966).

78 Gamble, T. & Zarkower, D. Identification of sex-specific molecular markers using restriction site associated DNA sequencing (RAD-seq). Molecular Ecology Resources, n/a-n/a, doi:10.1111/1755-0998.12237 (2014).

79 Palmer, D. H., Rogers, T. F., Dean, R. & Wright, A. E. How to identify sex chromosomes and their turnover. Mol Ecol 28, 4709–4724, doi:10.1111/mec.15245 (2019).

80 Waters, P. D., Duffy, B., Frost, C. J., Delbridge, M. L. & Graves, J. A. M. The human Y chromosome derives largely from a single autosomal region added to the sex chromosomes 80–130 million years ago. Cytogenetic and Genome Research 92, 74–79, doi:10.1159/000056872 (2001).

81 Prates, I. et al. Biogeographic links between southern Atlantic Forest and western South America: Rediscovery, re-description, and phylogenetic relationships of two rare montane anole lizards from Brazil. Mol Phylogenet Evol 113, 49–58, doi:10.1016/j.ympev.2017.05.009 (2017).

82 Velasco, J. A. et al. Climatic niche attributes and diversification in Anolis lizards. Journal of Biogeography 43, 134–144 (2016).

83 O’Leary, N. A. et al. Reference sequence (RefSeq) database at NCBI: current status, taxonomic expansion, and functional annotation. Nucleic Acids Res 44, D733–745, doi:10.1093/nar/gkv1189 (2016).

84 Bachtrog, D. Y-chromosome evolution: emerging insights into processes of Y-chromosome degeneration. Nature reviews. Genetics 14, 113–124, doi:10.1038/nrg3366 (2013).

85 Charlesworth, B., Coyne, J. & Barton, N. H. The relative rates of evolution of sex chromosomes and autosomes. American Naturalist 130, 113–146 (1987).

86 Singh, N. D., Larracuente, A. M. & Clark, A. G. Contrasting the Efficacy of Selection on the X and Autosomes in Drosophila. Molecular Biology and Evolution 25, 454–467, doi:10.1093/molbev/msm275 (2007).

87 Begun, D. J. & Whitley, P. Reduced X-linked nucleotide polymorphism in Drosophila simulans. Proceedings of the National Academy of Sciences of the United States of America 97, 5960–5965, doi:10.1073/pnas.97.11.5960 (2000).

88 Bachtrog, D., Jensen, J. D. & Zhang, Z. Accelerated adaptive evolution on a newly formed X chromosome. PLoS Biology 7, e1000082, doi:10.1371/journal.pbio (2009).

89 van de Schoot, M. Within and between island radiation and genetic variation in Anolis sagrei MSc thesis, Wageningen University, (2016).

90 Lieberman-Aiden, E. et al. Comprehensive mapping of long-range interactions reveals folding principles of the human genome. Science 326, 289–293, doi:10.1126/science.1181369 (2009).

91 Bolger, A. M., Lohse, M. & Usadel, B. Trimmomatic: a flexible trimmer for Illumina sequence data. Bioinformatics 30, 2114–2120, doi:10.1093/bioinformatics/btu170 (2014).

92 Paulino, D. et al. Sealer: a scalable gap-closing application for finishing draft genomes. BMC Bioinformatics 16, 230, doi:10.1186/s12859-015-0663-4 (2015).

93 Li, H. Aligning sequence reads, clone sequences and assembly contigs with BWA-MEM. 1303.3997 (2013).

94 Li, R. et al. De novo assembly of human genomes with massively parallel short read sequencing. Genome Research 20, 265–272, doi:papers3://publication/uuid/5D10FD6E-43D1-49C9-A0F9-4B5E618294A1 (2010).

95 Altschul, S. F., Gish, W., Miller, W., Myers, E. W. & Lipman, D. J. Basic local alignment search tool. J Mol Biol 215, 403–410, doi:10.1016/S0022-2836(05)80360-2 (1990).

96 Nakabayashi, R. & Morishita, S. HiC-Hiker: a probabilistic model to determine contig orientation in chromosome-length scaffolds with Hi-C. Bioinformatics 36, 3966–3974, doi:10.1093/bioinformatics/btaa288 (2020).

97 Mayjonade, B. et al. Extraction of high-molecular-weight genomic DNA for long-read sequencing of single molecules. Biotechniques 61, 203–205, doi:10.2144/000114460 (2016).

98 Hackl, T., Hedrich, R., Schultz, J. & Forster, F. proovread: large-scale high-accuracy PacBio correction through iterative short read consensus. Bioinformatics 30, 3004–3011, doi:10.1093/bioinformatics/btu392 (2014).

99 Wang, J. R., Holt, J., McMillan, L. & Jones, C. D. FMLRC: Hybrid long read error correction using an FM-index. BMC Bioinformatics 19, 50, doi:10.1186/s12859-018-2051-3 (2018).

100 Boetzer, M. & Pirovano, W. SSPACE-LongRead: scaffolding bacterial draft genomes using long read sequence information. BMC Bioinformatics 15, 211, doi:10.1186/1471-2105-15-211 (2014).

101 Pryszcz, L. P. & Gabaldón, T. Redundans: an assembly pipeline for highly heterozygous genomes. Nucleic Acids Research 44, e113–e113, doi:10.1093/nar/gkw294 (2016).

102 Xu, G. C. et al. LR_Gapcloser: a tiling path-based gap closer that uses long reads to complete genome assembly. Gigascience 8, doi:10.1093/gigascience/giy157 (2019).

103 Robinson, J. T. et al. Juicebox.js Provides a Cloud-Based Visualization System for Hi-C Data. Cell Syst 6, 256–258.e251, doi:10.1016/j.cels.2018.01.001 (2018).

104 Bernt, M. et al. MITOS: Improved de novo metazoan mitochondrial genome annotation. Molecular Phylogenetics and Evolution 69, 313–319, doi:https://doi.org/10.1016/j.ympev.2012.08.023 (2013).

105 R: A Language and Environment for Statistical Computing (R Foundation for Statistical Computing, Vienna, Austria, 2020).

106 RepeatModeler Open-1.0 (2008-2015).

107 RepeatMasker Open-4.0 (2013-2015).

108 Boer, E. F. et al. Pigeon foot feathering reveals conserved limb identity networks. Developmental Biology 454, 128–144, doi:https://doi.org/10.1016/j.ydbio.2019.06.015 (2019).

109 Akashi, H. D., Cadiz Diaz, A., Shigenobu, S., Makino, T. & Kawata, M. Differentially expressed genes associated with adaptation to different thermal environments in three sympatric Cuban Anolis lizards. Molecular Ecology 25, 2273–2285, doi:10.1111/mec.13625 (2016).

110 Trapnell, C. et al. Differential gene and transcript expression analysis of RNA-seq experiments with TopHat and Cufflinks. Nature Protocols 7, 562–578, doi:papers3://publication/doi/10.1038/nprot.2012.016 (2012).

111 Fu, L., Niu, B., Zhu, Z., Wu, S. & Li, W. CD-HIT: accelerated for clustering the next-generation sequencing data. Bioinformatics 28, 3150–3152, doi:10.1093/bioinformatics/bts565 (2012).

112 Li, W. & Godzik, A. Cd-hit: a fast program for clustering and comparing large sets of protein or nucleotide sequences. Bioinformatics 22, 1658–1659, doi:10.1093/bioinformatics/btl158 (2006).

113 Li, H. et al. The Sequence Alignment/Map format and SAMtools. Bioinformatics 25, 2078–2079, doi:10.1093/bioinformatics/btp352 (2009).

114 Li, H. Improving SNP discovery by base alignment quality. Bioinformatics 27, 1157–1158, doi:10.1093/bioinformatics/btr076 (2011).

115 Danecek, P. et al. The variant call format and VCFtools. Bioinformatics 27, 2156–2158, doi:10.1093/bioinformatics/btr330 (2011).

116 Quinlan, A. R. BEDTools: The Swiss-Army Tool for Genome Feature Analysis. Curr Protoc Bioinformatics 47, 11.12.11–34, doi:10.1002/0471250953.bi1112s47 (2014).

117 Cingolani, P. et al. A program for annotating and predicting the effects of single nucleotide polymorphisms, SnpEff: SNPs in the genome of Drosophila melanogaster strain w1118; iso-2; iso-3. Fly (Austin) 6, 80–92, doi:10.4161/fly.19695 (2012).

118 Grabherr, M. G. et al. Genome-wide synteny through highly sensitive sequence alignment: Satsuma. Bioinformatics 26, 1145–1151, doi:10.1093/bioinformatics/btq102 (2010).

119 Krzywinski, M. et al. Circos: an information aesthetic for comparative genomics. Genome research 19, 1639–1645 (2009).

120 Hao, Z. et al. RIdeogram: drawing SVG graphics to visualize and map genome-wide data on the idiograms. PeerJ Computer Science 6, e251 (2020).

121 Kichigin, I. G. et al. Evolutionary dynamics of Anolis sex chromosomes revealed by sequencing of flow sorting-derived microchromosome-specific DNA. Mol Genet Genomics 291, 1955–1966, doi:10.1007/s00438-016-1230-z (2016).

122 Peterson, B. K., Weber, J. N., Kay, E. H., Fisher, H. S. & Hoekstra, H. E. Double Digest RADseq: An Inexpensive Method for De Novo SNP Discovery and Genotyping in Model and Non-Model Species. PLoS ONE 7, e37135, doi:10.1371/journal.pone.0037135.t001 (2012).

123 Eaton, D. A. PyRAD: assembly of de novo RADseq loci for phylogenetic analyses. Bioinformatics 30, 1844–1849 (2014).

124 Puritz, J. B., Hollenbeck, C. M. & Gold, J. R. dDocent: a RADseq, variant-calling pipeline designed for population genomics of non-model organisms. PeerJ 2, e431, doi:10.7717/peerj.431 (2014).

125 Li, H. & Durbin, R. Fast and accurate short read alignment with Burrows-Wheeler transform. Bioinformatics 25, 1754–1760, doi:10.1093/bioinformatics/btp324 (2009).

126 Garrison, E. & Marth, G. Haplotype-based variant detection from short-read sequencing. arXiv preprint 1207.3907 (2012).

127 Purcell, S. et al. PLINK: A Tool Set for Whole-Genome Association and Population-Based Linkage Analyses. The American Journal of Human Genetics 81, 559–575, doi:10.1086/519795 (2007).

128 Browning, B. L. & Browning, S. R. Genotype Imputation with Millions of Reference Samples. Am J Hum Genet 98, 116–126, doi:10.1016/j.ajhg.2015.11.020 (2016).

