## Supplementary Figures and Methods for "Chromosome-scale genome assembly of the brown anole (*Anolis sagrei*), a model species for evolution and ecology"

### Supplementary Methods

#### Publication record

We performed separate queries for "*Anolis sagrei*" and "*Anolis carolinensis*" appearing in the title or abstract of publications using the http://dimensions.ai database. We limited the query to the years 1980-2000 inclusively. These query results were merged and the number of publications per species per year were calculated in R v3.6.3[1] and plotted using ggplot2 v 3.3.3[2].

### Supplementary References

1 R: A Language and Environment for Statistical Computing (R Foundation for Statistical Computing, Vienna, Austria, 2020).

2 Wickham, H. *ggplot2: Elegant Graphics for Data Analysis*. (Springer International Publishing, 2016).

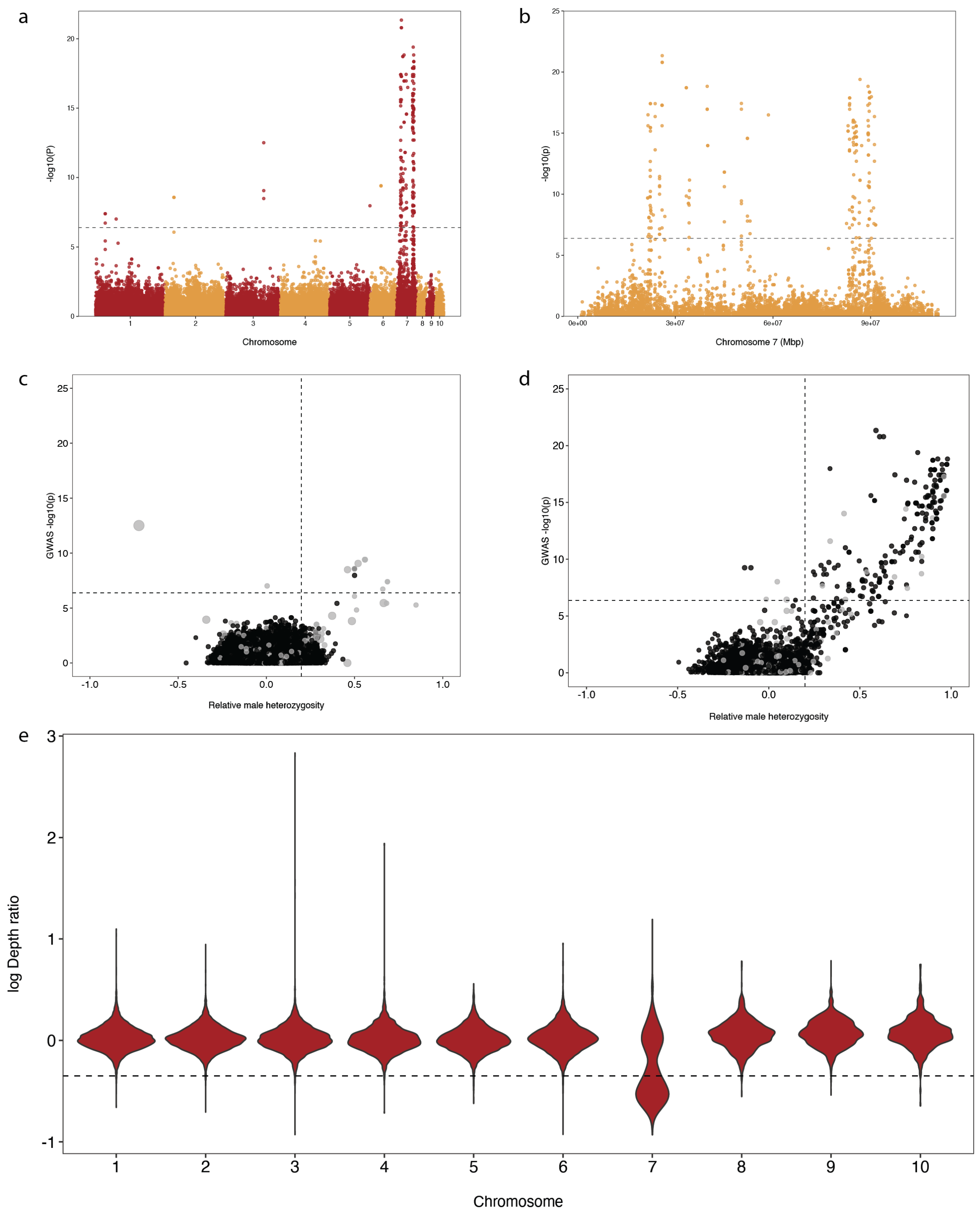


**Supplementary Figure 1**: **Identification of the sex chromosome of *Anolis sagrei*. a**, Manhattan plot for the sex GWAS analysis, showing only the largest 10 scaffolds. The dotted line represents the 5% significance threshold after Bonferroni correction. **b**, Magnification of GWAS results for scaffold 7. **c,d**, Relative male heterozygosity, GWAS -log10 *P*-values, and male/female depth ratio for scaffolds 1-7 (**c**) and for scaffold 7 (**d**). The horizontal dotted line is the GWAS significance threshold, and the vertical line is the threshold for the identification of heterozygosity outliers. SNPs with higher depth in males than in females (i.e. genomic outliers for depth ratio) are shown in grey. Circle size is proportional to depth ratio. **e**, Log-transformed male/female depth ratio for scaffolds 1-10. The dotted line marks the threshold used for the classification of genomic outliers with low depth ratio.

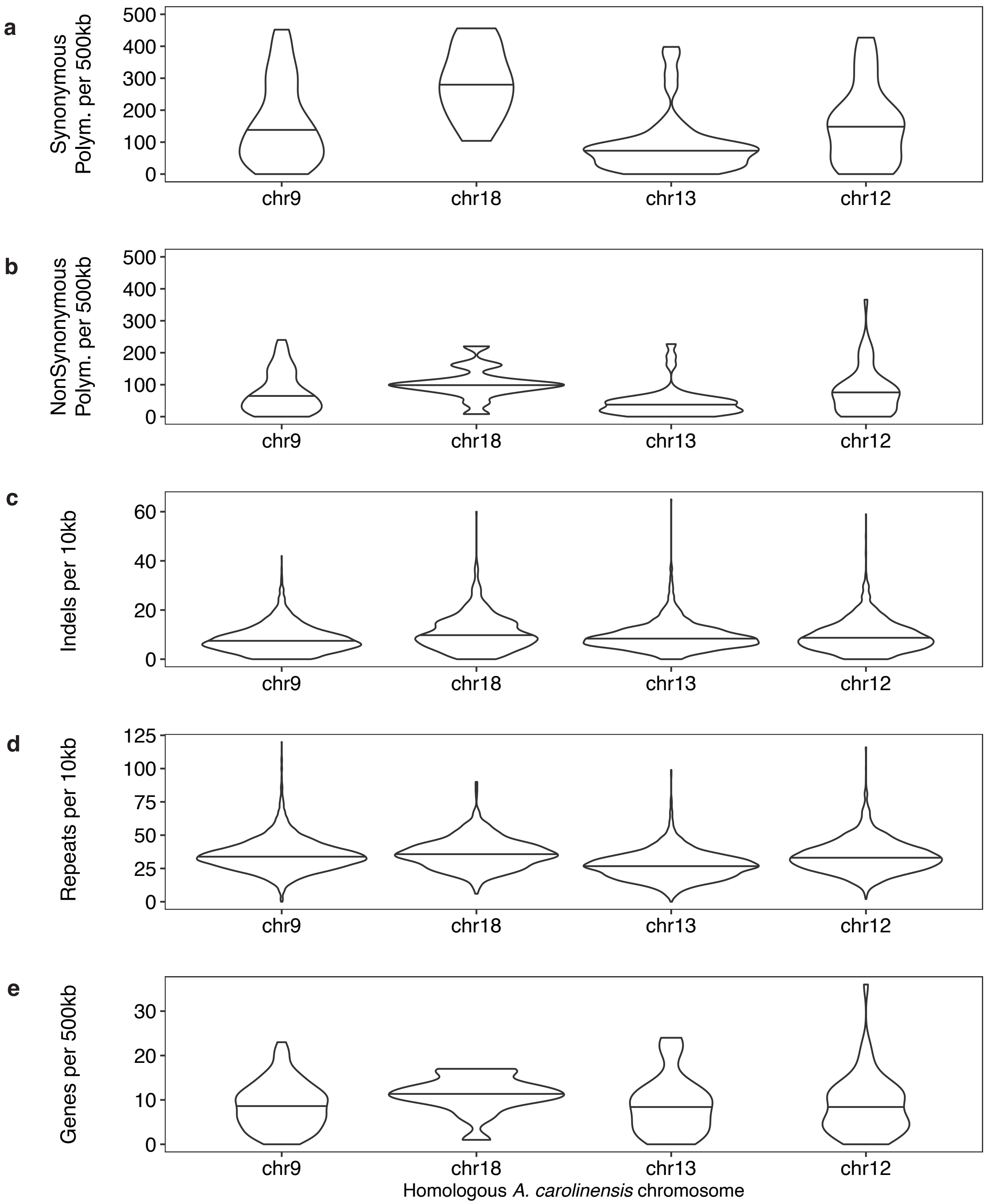


**Supplementary Figure S2. Differences among *Anolis sagrei* X chromosome sub compartments.** Regions of the *Anolis sagrei* X chromosome homologous to autosomes in *A. carolinensis* have greater densities **a**) synonymous polymorphisms, **b**) nonsynonymous polymorphisms, **c**) indels, **d**) repetitive genetic elements, and **e**) genes relative to the region homologous to the *A. carolinensis* X (chromosome 13).

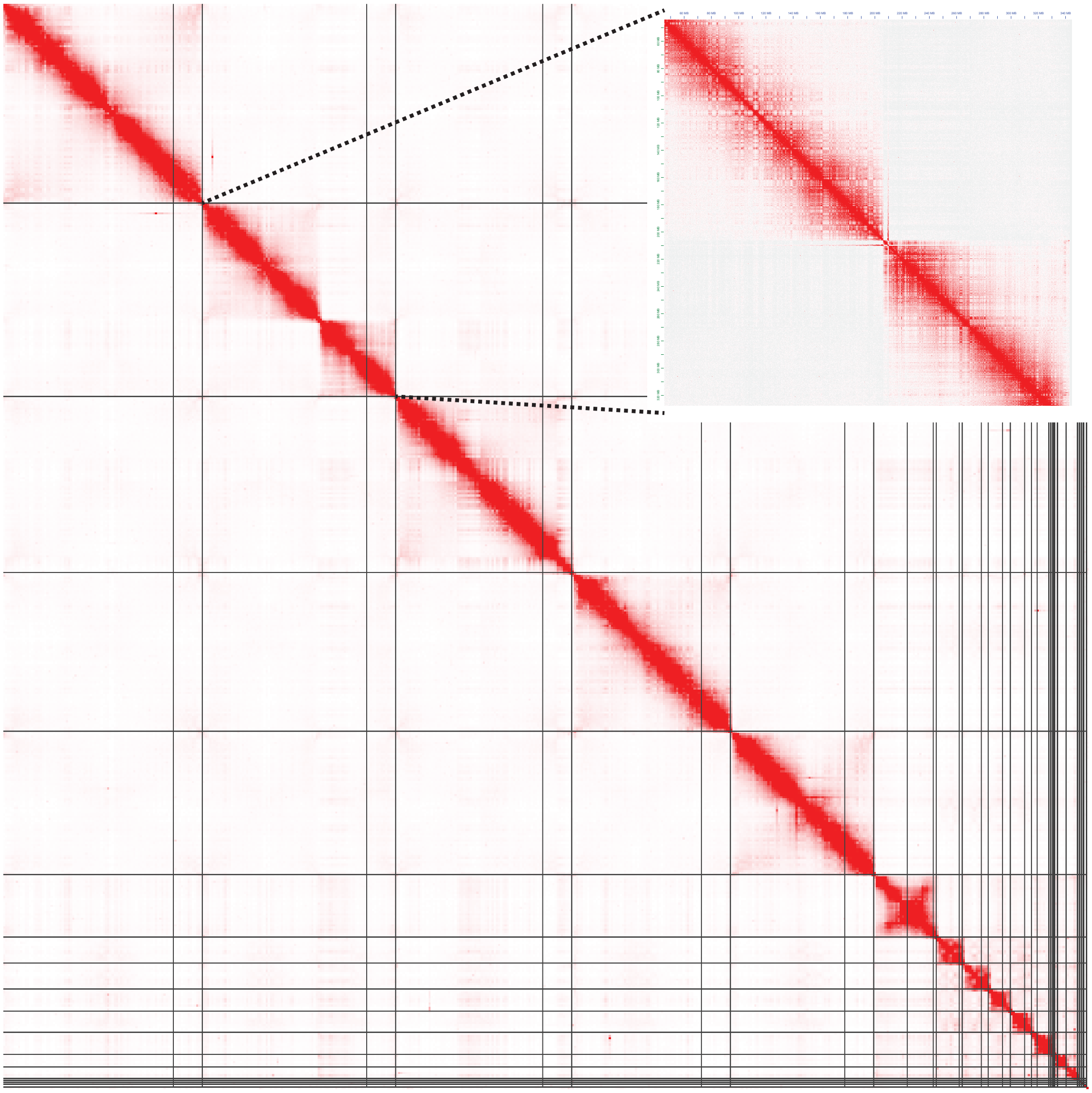

**Supplementary Figure S3. HiC Link Density Histogram of *Anolis sagrei* assembly version 2.0.** Scaffold 2 (enlarged in the inset) of this assembly is an assembly mis-join of two smaller chromosomes based on the lack of HiC read pairs mapping across center of this scaffold.
